# Brain-wide projections of mouse dopaminergic zona incerta neurons

**DOI:** 10.1101/2024.09.06.611701

**Authors:** Bianca S. Bono, Kenichiro Negishi, Yasmina Dumiaty, Monica S. Ponce, Titilayo C. Akinbode, Kayla S. Baker, C. Duncan P. Spencer, Elizabeth Mejia, Marina Guirguis, Alex J. Hebert, Arshad M. Khan, Melissa J. Chee

## Abstract

The zona incerta (ZI) supports diverse behaviors including binge feeding, sleep/wake cycles, nociception, and hunting. This diversity of functions can be attributed to the heterogenous neurochemicals, cytoarchitecture, and efferent connections that characterize the ZI. The ZI is predominantly GABAergic, but we recently identified a subset of medial ZI GABA cells that co-express dopamine (DA), as marked by the enzyme tyrosine hydroxylase (TH). While the role of GABA within the ZI is well studied, little is understood about the function of ZI DA cells. To identify potential roles of ZI DA cells we mapped the efferent fiber projections from *Th-cre* ZI cells. We first validated a *Th-cre;L10-Egfp* mouse line and found that medial *Egfp* ZI cells were more likely to co-express TH-immunoreactivity (TH-ir). We thus delivered a cre-dependent virus into the medial ZI of *Th-cre* or *Th-cre;L10-Egfp* mice and selected two injection cases for full brain mapping. We selected the cases with the lowest (17%) and highest (53%) percentage of colocalization between TH-ir and virus transfected cells labelled with DsRed. Overall, DsRed-labelled fibers were observed throughout the brain and were most prominent within motor-related regions of the midbrain (MBmot), notably the periaqueductal grey area and superior colliculus. We also observed considerable DsRed-labelled fibers within the polymodal cortex associated regions of the thalamus (DORpm), including the paraventricular thalamic nucleus and nucleus of reunions. Overall, ZI DA cells displayed a similar connectivity profile to ZI GABA cells, suggesting that ZI DA cells may perform synergistic or opposing functions at the same target sites.

**List of RRIDs:** AB_2201528, AB_10013483, AB_11177031, AB_2340593, AB_2315778, SCR_016477

**Three key points:**

- Tyrosine hydroxylase immunoreactivity was more prominent in the medial zona incerta
- Intersection points from two *Th-cre* injection cases revealed common target regions
- Dopaminergic zona incerta cells project predominantly to motor-related brain regions

## 1 INTRODUCTION

The zona incerta (ZI) is situated in the subthalamic region between the thalamus and hypothalamus and first captured scientific interest over a century ago because of its strategic position to several important fiber systems in the human brain (Forel, 1877; Malone, 1910; Nauta and Haymaker 1969). Comparative neuroanatomical studies (Ariëns Kappers et al., 1965) have since supported the view that neuronal populations in the ZI are associated with prominent fiber systems that traverse its gray matter expanse. Cajal (1911/1995) provided the first detailed drawings of mouse ZI neurons from Golgi-stained material, and these drawings identified ascending medial lemniscal, bidirectional internal capsular, and descending corticothalamic fiber systems in close association with ZI cell populations. Together, these early works surmised that the ZI plays important roles in sensorimotor communication.

Consistent with initial observations by Cajal are characterizations of the ZI as a subthalamic structure that links striatal structures with midbrain tegmental structures (Le Gros Clark, 1938). However, the ZI can also be considered part of the ’extrapyramidal’ motor system involving an underlying hypothalamic element (Kuhlenbeck, 1977). Indeed, the ZI is currently being examined as a clinical target for movement disorders like Parkinson’s disease (Blomstedt et al., 2018) as deep brain stimulation of the caudal ZI reduces muscle rigidity and tremors (Plaha et al., 2006, 2008). On the sensory side, functional neuroanatomical work in the ZI of rats have identified it as an important paralemniscal pathway conveying somatosensory information to the cortex for vibrissal sensations, in a pathway set apart from medial lemniscal pathways making first-order contact with sensory thalamic structures (Diamond and Ahissar, 2007; Urbain and Deschênes, 2007). In addition to somatosensation, the ZI also integrates input from varied sensory modalities, including pain (Masri et al., 2009; Moon and Park, 2017), visual (Zhao et al., 2019), olfactory (Li et al., 2024), and auditory input (Shang et al., 2019; Li et al., 2024). Collectively, neuroanatomical and functional studies have defined the ZI as an important structure in sensorimotor processing and motor coordination, and the ZI is recognized as an integrative node that permits the organism to interact and respond to its environment (Wang et al., 2020b). For example, optogenetic activation of the ZI is important for the integration of somatic, visual, and auditory functions during hunting (Zhao et al., 2019), and activating the ZI can suppress flight responses from a loud sound or suppress defensive behaviors (Chou et al., 2018).

Our understanding of the ZI has been refined by chemoarchitectural characterization of ZI cell types across various taxa. In mouse and other taxa, the ZI is predominantly GABAergic (Lin et al., 1990; Yang et al., 2022), but the ZI is also a heterogeneous structure that produce other markers or chemical messengers, including glutamate (Beitz, 1989; Border and Mihailoff, 1991), serotonin (Bosler et al., 1984), catecholamines (Chan-Palay et al., 1984), calbindin (Celio, 1990), and parvalbumin. However, ZI GABA cells may also co-express chemical messengers and markers such as somatostatin (Finley et al., 1981; Lin et al., 2023) and the LIM homeobox 6 (Lhx6) transcription factor (Liu et al., 2017). The heterogeneity of the ZI may help explain its role in diverse behavioral and physiological functions. In addition to defensive behaviors (Chou et al., 2018; Lin et al., 2023) and hunting (Zhao et al., 2019), the ZI has also been implicated in feeding (Zhang and van den Pol, 2017; Ye et al.), the regulation of sleep and wake cycles (Liu et al., 2017), the suppression of fear generalization (Venkataraman et al., 2019) and nociception (Singh et al., 2022). The functional diversity of the ZI implicates that it has numerous target sites, and a recent study has confirmed that GABAergic ZI cells has widespread, brain-wide projections (Yang et al., 2022). However, it is not yet known if each specific ZI cell type contributes similarly or uniquely to the overall distribution of ZI projections.

We recently showed that a subset of GABAergic ZI cells that were marked by the co-expression of tyrosine hydroxylase (TH) also contained dopamine (DA) (Negishi et al., 2020). This work suggested that this group of ZI cells formed a novel subpopulation that could release both GABA and DA. However, the specific targets of these GABAergic ZI DA neurons are not yet known. In this study, we used a *Th-cre* transgenic mouse line to trace and map the fiber projections of ZI DA cells using an anterograde, Cre-dependent viral tracer. We overlaid the pattern of fiber distribution from two injection cases to delineate common projection targets, and we found that this GABAergic ZI DA population also projects widely throughout the brain. The distribution of *Th-cre* fibers overlapped with known targets of ZI neurons (Yang et al., 2022) but within a refined subset of those targets, including notable fibers within motor-related regions of the midbrain, such as the periaqueductal grey area and superior colliculus, as well as polymodal regions of the thalamus such as the paraventricular nucleus of the thalamus. Our findings suggest that GABAergic ZI DA cells may contribute to many behaviours previously ascribed to ZI GABA cells, especially hunting, defensive behaviors, and binge-like eating.

## 2 MATERIALS AND METHODS

### 2.1 Subjects

All experimental procedures were performed in accordance with the Animal Care Committee at Carleton University. Mice were group-housed with a 12:12 hour light-dark cycle (21–22 °C) and provided water and standard rodent chow *ad libitum* (2014 Teklad, Envigo, Mississauga, Canada). To visualize TH neurons, the *Th-IRES-cre* (Lindeberg et al., 2004) mouse was crossed to a *R26*-*lox-STOP-lox-L10-Egfp* reporter mouse (Krashes et al., 2014) to produce *Th-cre;L10-Egfp* mice expressing enhanced green fluorescent protein (EGFP) under the *Th* promoter.

### 2.2 Antibody characterization

The primary antibodies used for immunohistochemistry (IHC) are listed in **Table 1**.

**Table 1.** Primary antibodies used for immunohistochemistry.

| Antibody | Immunogen | Clonality,<br>Isotype | Source,<br>Catalog no.,<br>Lot no. | RRID | Titer |
| --- | --- | --- | --- | --- | --- |
| Mouse anti-tyrosine hydroxylase | Purified from PC12 cells | Monoclonal, IgG | MilliporeSigma, MAB318, 2849094 | AB_2201528 | 1:2,000 |
| Rabbit anti-dsRed | Full-length dsRed-Express | Polyclonal, IgG | Takara Bio USA, 632496, 1202020 | AB_10013483 | 1:1,000 |
| Sheep anti-tyrosine hydroxylase | Purified from pheochromocytoma | Polyclonal, IgG | GeneTex, GTX82570, 822005648 | AB_11177031 | 1:1,000 |

#### Rabbit anti-DsRed antibody

was raised against a variant of the *Discosoma sp*. red fluorescent protein and can label various red fluorescent proteins, including mCherry and tdTomato. Antibody specificity was demonstrated by the absence of immunostaining in cre-expressing tissues that do not produce tdTomato (Cheng et al., 2009; Chee et al., 2013).

#### Mouse anti-TH antibody

was produced by purifying PC12 cells. Specificity of this antibody was determined by unilaterally lesioning the ventral tegmental area and substantia nigra in rats (Chung et al., 2008; Bourdy et al., 2014). Further specificity of this antibody has been shown in zebrafish, whereby knocking out *Th1*, a gene that encodes TH in the central nervous system of zebrafish, abolishes TH-ir in the brain (Kuscha et al., 2012).

#### Sheep anti-TH antibody

was produced by purification from pheochromocytomas. Specificity of this antibody was determined by lower TH-expressing cells in the substantia nigra following unilateral, 6-hydroxydopamine hydrobromide lesion to the rat striatum of rats (Choudhury et al., 2011).

The secondary antibodies used are listed in **Table 2** and were raised in a donkey against the species of the corresponding primary antibody (i.e., rabbit, mouse, sheep).

**Table 2.** Secondary antibodies used for immunohistochemistry.

| Antibody | Immunogen | Source,<br>Catalog no.,<br>Lot no. | RRID | Titer |
| --- | --- | --- | --- | --- |
| Donkey anti-mouse DyLight 649 | IgG (H+L) | Jackson ImmunoResearch, 715-496-150<sup>a</sup>, 82079 | n/a<sup>a</sup> | 1:500 |
| Biotin-conjugated donkey anti-rabbit | IgG (H+L) | Jackson ImmunoResearch, 711-065-152, 116529 | AB_2340593 | 1:1,000 |
| Donkey anti-sheep Cy 3 | IgG (H+L) | Jackson ImmunoResearch, 713-165-147, 148268 | AB_2315778 | 1:500 |

<sup>a</sup> This item has since been discontinued; no RRID is available

### 2.3 Stereotaxic injections

Male (4–8 wk old) *Th-cre* or *Th-cre;L10-Egfp* mice (N = 7) were administered meloxicam (5 mg/kg) analgesia subcutaneously (sc) and then secured in a stereotaxic frame (Kopf Instruments, Tujunga, CA) under deep isoflurane anesthesia. Mice were unilaterally injected with 25–75 nl (25 nl/min) of a Cre-dependent adeno-associated virus (AAV) encoding mCherry (AAV8–EF1α–DIO–ChR2(H134R)–mCherry; 4.4 × 10^12^ genomic copies/ml, lot AAV2037, Canadian Neurophotonics Platform Viral Vector Core Facility [RRID: SCR_016477]) or tdTomato (AAV2/DJ–EF1α–DIO–ChETA–tdTomato; 1.7 × 10^13^ genomic copies/ml, lot AAV1131, Canadian Neurophotonics Platform) at the medial ZI using coordinates (in mm) relative to Bregma at the skull surface: anteroposterior, −1.40 or −1.50; mediolateral, −0.50 or −0.70; dorsoventral, −4.70. Mice were allowed to recover for four weeks to promote viral transduction and reporter expression along nerve terminals.

### 2.4 Tissue processing

Brain tissue was prepared and collected as previously described (Bono et al., 2022), unless indicated otherwise, for each procedure below.

#### 2.4.1 *In situ* hybridization (ISH)

Brain tissue from male *Th-cre;L10-Egfp* mice (n = 3, 8–10 wk old) was collected in phosphate buffered saline (PBS) and immediately mounted onto Superfrost Plus microscope slides (Fisher Scientific, Hampton, NH). The slides were left to air-dry at room temperature (RT; 21–22 °C) for one hour, at −20 °C overnight, and then packed into slide boxes for storage at −80 °C.

#### 2.4.2 Injection case validation and mapping

Brain tissue from male *Th-cre* or *Th-cre;L10-Egfp* mice was sliced into four series of 20–30 μm-thick sections and stored at −80 °C in an antifreeze solution comprising 50% formalin, 20% glycerol, and 30% ethylene glycol. Tissues were later processed for indirect fluorescence or immunoperoxidase IHC.

### 2.5 Nissl staining

Brain tissues underwent Nissl stain as previously described (Negishi et al., 2020; Bono et al., 2022; Miller et al., 2024).

### 2.6 Dual-label ISH and IHC

*Th-cre;L10-Egfp* brain tissues underwent target retrieval and Protease III treatment as previously described (Bono et al., 2022). In brief, RNAscope probes (**Table 3**) *Mm-Ppib* for positive control targeting, *Bacillus dapB* for negative control targeting, or *Egfp-O2* and *Mm-Th* for experimental targeting were added to their respective slides and hybridized for 2 h at 40 °C. *Egfp* and *Th* probes were added to probe diluent (1:50; 300041, ACD) as neither probe occupied Channel 1. Following AMP-mediated amplification, the tissues were first treated with HRP-C2 (15 min, 40 °C; 323105, ACD) to develop *Egfp* hybridization visualized with Opal 520 (FP1487001KT, PerkinElmer, Waltham, MA) diluted in TSA buffer (1:750; 322809, ACD) and then with HRP-C3 (15 min, 40 °C; 323106, ACD) to develop *Th* hybridization visualized with Opal 690 (FP1497001KT, Perkin Elmer) diluted in TSA buffer (1:750).

**Table 3.**
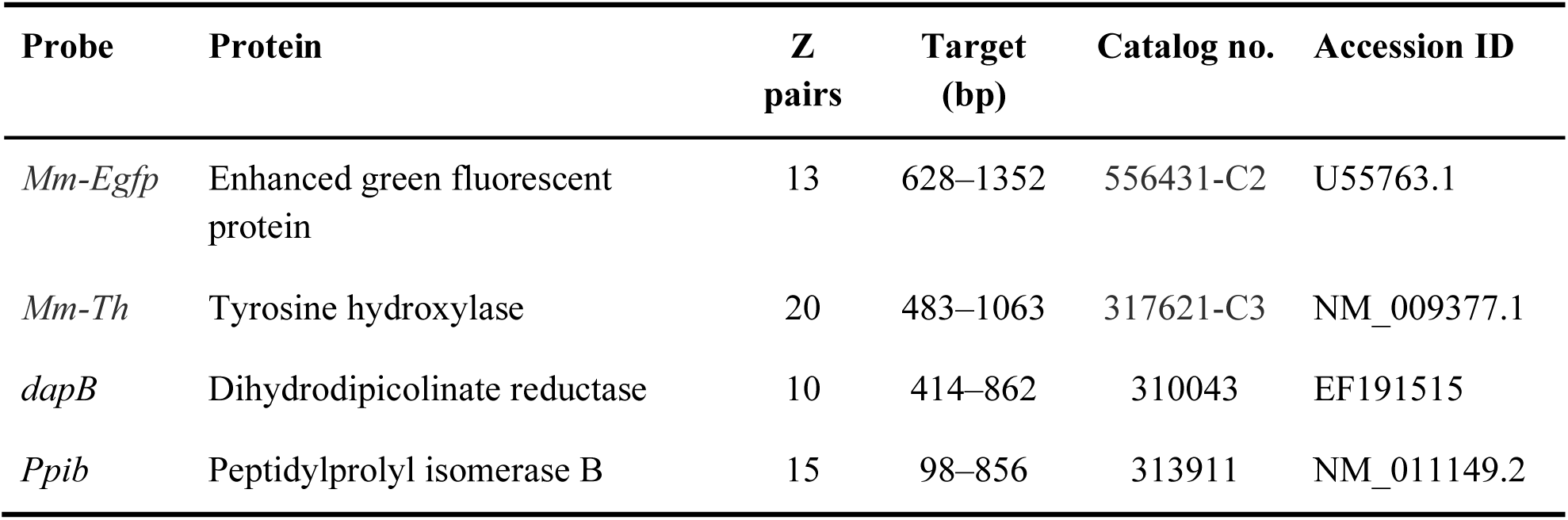
List of RNAscope probes for *in situ* hybridization.

After ISH treatment, tissues were processed for IHC as previously described (Bono et al., 2022). In brief, the tissues were incubated with a sheep anti-TH (1:1,000; diluted in PBS) for 30 min at room temperature (RT, 23 °C) and then a donkey anti-sheep Cyanine (Cy) 3 (1:500; diluted in PBS) for 30 min (RT).

### 2.7 Indirect fluorescence IHC

All injection cases underwent IHC processing as previously described (Shammah-Lagnado et al., 1985; Chee et al., 2013; Tapia et al., 2023) to determine viral spread at the injection site and the extent of viral transduction in TH-immunoreactive (-ir) cells. In brief, free-floating brain sections were incubated with a rabbit anti-DsRed (1:1,000) and mouse anti-TH (1:2,000) primary antibodies for 12 h (RT) and then a donkey anti-rabbit Cy 3 (1:500) and donkey anti-mouse DyLight 649 (1:500) secondary antibodies for 2 h (RT).

### 2.8 Indirect immunoperoxidase IHC

As previously described (Chee et al., 2013; Tapia et al., 2023), injection cases selected for fiber mapping underwent IHC processing with an immunoperoxidase reaction to enhance labeling of virally-transduced *Th-cre* fiber projections. In brief, free-floating brain sections were blocked in 3% normal donkey serum for 2 h (Jackson ImmunoResearch Laboratories, Inc., West Grove, PA) and then incubated with rabbit anti-DsRed primary antibody (1:1,000) for 24–48 h at 4 °C and then a biotin-conjugated donkey anti-rabbit secondary antibody (1:1,000) for 5 h (RT). After rinsing in five 5-min exchanges of Tris-buffered saline (TBS; RT), the sections were treated to form avidin-biotin horseradish peroxidase complexes (ABC) at biotin-conjugated sites using the Vectastain Elite ABC kit (0.45% v/v Reagent A and B in TBS containing 0.1% Triton X-100; PK-6100, Vector Laboratories) for 1 h (RT). Ds-Red immunoreactivity was then developed by reacting the sections in a 0.05% diaminobenzidine and 0.015% hydrogen peroxide TBS cocktail for 15–20 min (RT). Finally, the sections were washed in five 5-min exchanges of TBS and mounted onto gelatin coated slides. After air-drying, the slides were dehydrated in ascending concentrations of ethanol (50%, 70%, 95%, and 100% ×3 for 3 min each), delipidated in xylene (25 min), and coverslipped with DPX mountant (06522, Sigma Aldrich, St. Louis, MO).

### 2.9 Microscopy

#### 2.9.1 Brightfield imaging

Large field-of-view photomicrographs of Nissl-stained brain sections were acquired via a Plan Apochromat ×4 objective lens (0.20 numerical aperture) mounted on a Nikon Eclipse Ti2 inverted microscope (Nikon Instruments Inc., Mississauga, Canada) equipped with a fully motorized stage and DS-Ri2 color camera (Nikon) as previously described (Negishi et al., 2020; Bono et al., 2022). High magnification photomicrographs to differentiate between axon fibers and terminals were acquired via a ×40 objective lens (0.95 numerical aperture)

#### 2.9.2 Epifluorescence imaging

Large field-of-view photomicrographs of DAPI-stained nuclei or DsRed-ir cells were acquired using a Prime 95B CMOS camera (Photometrics, Tucson, AZ). DAPI- or Cy 3-labelled excitation was provided by a SPECTRA X light engine (Lumencor, Beaverton, OR) passing through a ET395/25x or ET550/15x excitation filter (Chroma Technology Corporation, Bellows Falls, VT), respectively.

#### 2.9.3 Confocal imaging

High magnification photomicrographs were generated using a Nikon C2 confocal system fitted with a Plan Apochromat ×10 objective lens (0.45 numerical aperture). Regions of interest were identified by epifluorescence imaging (as described in 2.9.2).

To analyze ISH-treated tissue, photomicrographs were generated with 405-, 488-, 561-, and 640-nm wavelength lasers to visualize DAPI-, Opal 520-, Opal 690-, and Cy 3-labeled signals, which were pseudo-colored dark blue, green, magenta, and light blue, respectively. The same image acquisition settings were used to acquire images from experimental tissue, *dapB*-treated negative control tissue, and *Ppib*-treated positive control tissue. Photomicrographs from experimental tissue were adjusted using the same image processing settings applied to photomicrographs captured from *dapB*-hybridized tissues until no fluorescence was visible (Bono et al., 2022); this subtracted the background or fluorescence arising from non-specific binding.

To analyze the spread of injection sites, photomicrographs were generated with 561- and 640-nm wavelength lasers to visualize Cy 3- and DyLight 649-labeled signals, which were pseudo-colored red and light blue respectively.

#### 2.9.4 Darkfield imaging

Wide-field photomicrographs of DAB-stained sections were acquired with a ×10 objective lens (0.40 numerical aperture) mounted on an Olympus BX-63 microscope (Olympus Corporation, Tokyo, Japan) fitted with a darkfield condenser, motorized stage, and DP74 color camera. Tiled images were stitched and adjusted with cellSens Dimension imaging software (Version 2.3, Olympus) and exported as TIFF files.

### 2.10 Image and fiber analysis

All photomicrographs generated were tiled and stitched using NIS-Elements (Nikon Corporation, Konan, Japan), unless otherwise indicated. Any adjustments to the brightness, intensity, or contrast were applied to the entire photomicrograph via NIS-Elements and then the photomicrographs were exported as TIFF files in Illustrator 2023 (Adobe Inc., San Jose, CA). Any additional text, labels, or symbols (e.g., arrowheads or outlines) were added in Illustrator 2023.

#### 2.10.1 Plane-of-section analysis

Nissl-based parcellations were determined as previously described (Negishi et al., 2020; Bono et al., 2022).

#### 2.10.2 Atlas-based mapping

Nissl-based parcellations were transferred to large field-of-view epifluorescence images, which were aligned to their corresponding Nissl photomicrographs. The corresponding high magnification confocal photomicrographs were then aligned to the epifluorescence photomicrographs for either cell counting or fiber tracing. Cell bodies were labeled using the *Paintbrush* tool, and fibers were traced using the *Pen* tool (Illustrator 2023, Adobe). The labeled cell bodies or traced fibers were then transferred onto the corresponding *ARA* template (Dong, 2008). Differences in plane-of-section mediolaterally or dorsoventrally within a brain slice were carefully assessed so that angular differences in any dimension prompted a fragmented approach when mapping onto atlas templates (Simmons and Swanson, 2009).

#### 2.10.3 Quantification of ISH cells

Both *Egfp* and *Th* hybridization appeared as clusters of punctate “dots,” and *Egfp* or *Th* hybridized cells were marked using the Blob Brush tool in Illustrator if it expressed three or more dots per cell and if these dots localized to a DAPI-labelled soma. Only cell counts at *ARA* levels (L) that were available from at least two brains were included in our dataset. As previously described, cell counts were corrected for oversampling using the Abercrombie formula (Negishi et al., 2020; Bono et al., 2022), where mean tissue thickness (25.31 μm) was determined from 15 hypothalamic brain slices and mean cell diameter (12.82 μm) was determined from 50 cells within the ZI.

#### 2.10.4 Quantification of DsRed-labeled fibers

Each brain region was grouped into one of twenty subdivisions that we formed based on the hierarchical organization scheme (Wang et al., 2020a) and ontology (Allen Institute for Brain Science, 2011) provided by the *ARA* (**Supplemental Figure 1**). Large-scale quantification of DsRed-labeled fibers were determined from manual fiber tracings that were mapped onto *ARA* templates and then analyzed by each major brain division, which included: isocortex; olfactory regions of the cortex; hippocampal formation; cortical subplate of the cerebral cortex; striatum; palladium; sensory-motor and polymodal cortex related regions of the thalamus; periventricular zone, periventricular region, medial zone, and lateral zone of the hypothalamus; sensory, motor, and behavioral-state related regions of the midbrain; and sensory-motor, and behavioral-state related regions of the pons. Mapped fiber tracings from each major brain division were transferred to Photoshop 2023 (Adobe) to determine the density of fibers in the whole brain division per *ARA* level.

**Supplemental Figure 1.**
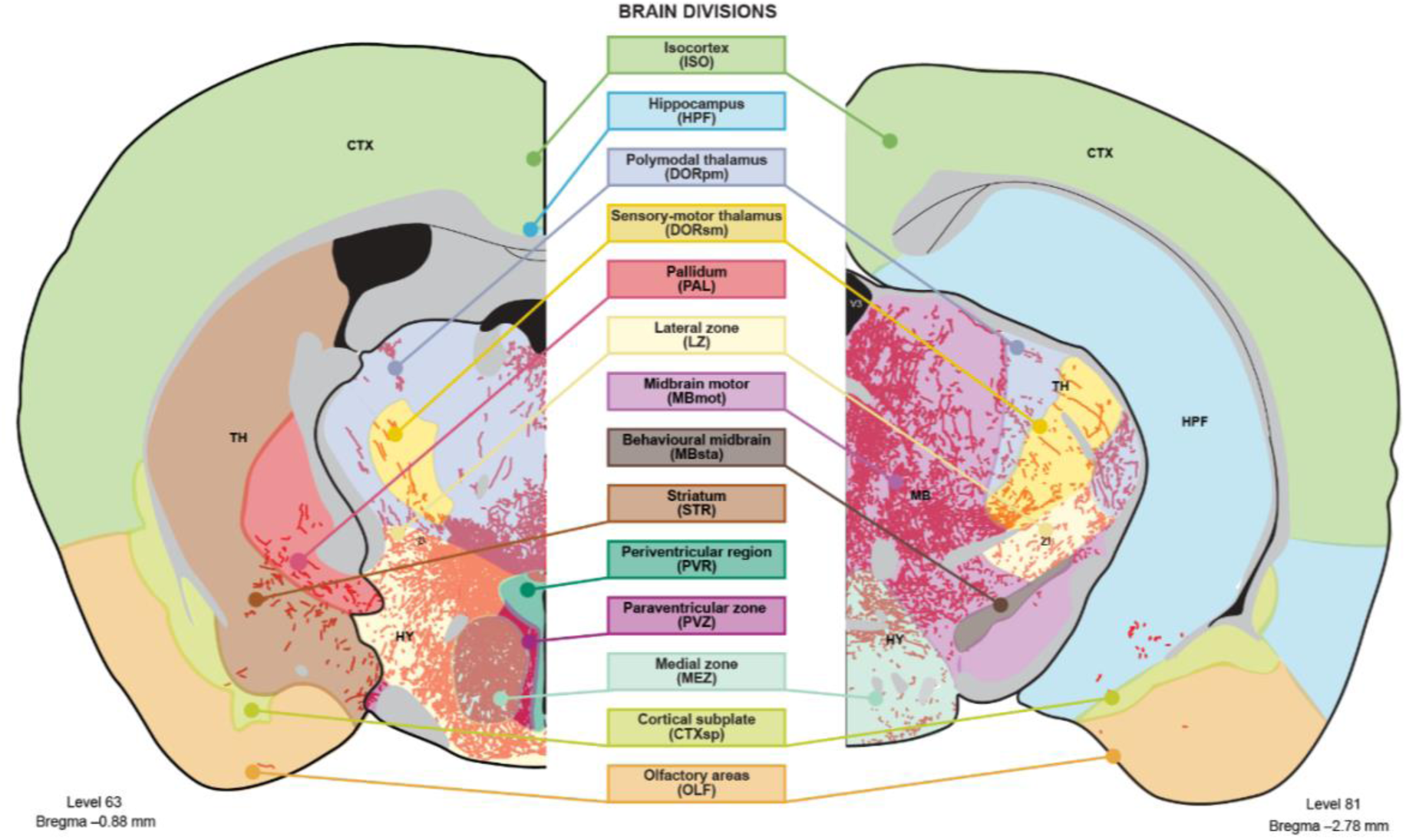
Hierarchical organization of major brain divisions. Mapped DsRed-labelled fibers were overlaid onto *Allen Reference Atlas* (Dong, 2008) templates. DsRed-labelled fibers were quantified based on their expression within each major brain division, as defined by the Allen Institute (Wang et al., 2020a; Allen Institute for Brain Science, 2011).

Fiber coverage was determined in each region of interest by first using the *lasso* tool to outline the borders of the region. Using the *Histogram* panel, we then selected “entire image” under the *source* dropdown menu to obtain the total number of pixels within the outlined region; this determined the total region area. Next, we adjusted the source menu to “selected layer” that contained the traced fibers to obtain the number of pixels occupied by a traced fiber.

#### 2.10.5 Intersecting fibers maps

Fiber maps from two injection cases were overlaid in separate layers using Illustrator 2023 (Adobe). The traced fibers from each injection case were converted into compound paths, and the compound paths from each injection case were overlaid upon one another. Compound paths that overlapped and mapped to the same region in both injection cases were identified using the *Intersect Pathfinder* effect in the *Pathfinder panel*.

### 2.11 Statistics

Statistical analyses and accompanying graphs were generated using GraphPad Prism 7 (GraphPad Software Inc., San Diego, CA). Pearson correlation coefficient with 95% confidence intervals was performed to determine the extent of correlation between fiber coverage between injection cases, as well as between the intersecting fiber coverage and the total region area. Likewise, a two-way ANOVA with Tukey’s HSD was performed to identify differences in the percentage of EGFP cell types within the medial and lateral ZI. Statistical significance was determined at p < 0.05. All data are presented as mean ± standard error of the mean (SEM).

## 3 RESULTS

### 3.1 Validation of *Th-cre* expression in the ZI

We evaluated the efficacy and specificity of Cre-mediated expression at the ZI of *Th-cre;L10-Egfp* mice (N = 3) by determining the colocalization of *Th* mRNA and TH-immunoreactivity in cells expressing *Egfp* mRNA. *Egfp* hybridized cells in the ZI spanned levels (L) 60–71 of the *Allen Reference Atlas* (*ARA*; Dong, 2008), but they were most abundant at L67–69 (**Figure 1a**).

**Figure 1.**
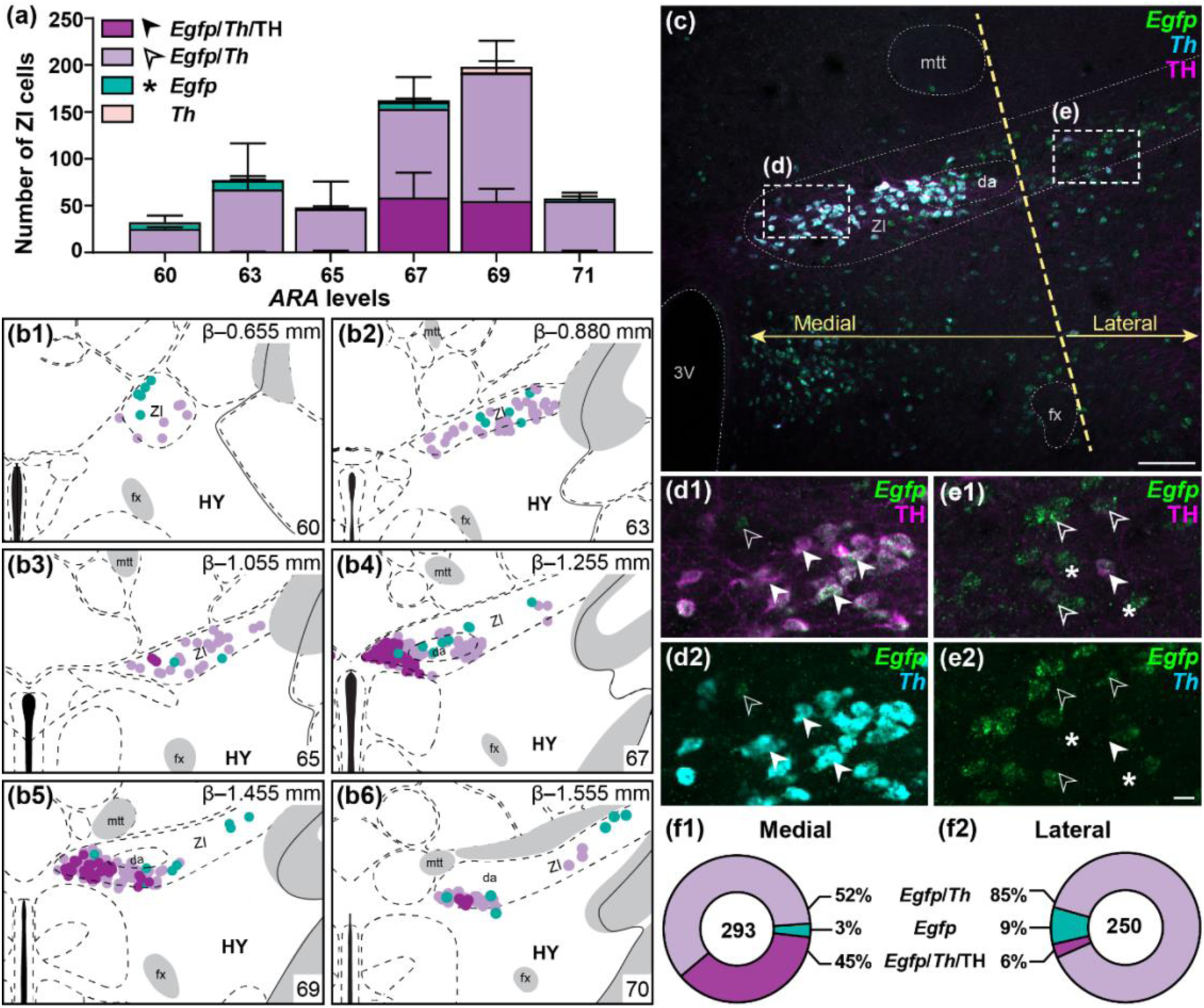
Robust colocalization of *Egfp* hybridization, *Th* hybridization, and TH immunoreactivity in the medial ZI. Number (**a**) and distribution of *Egfp* mRNA cells (**b**) that were ectopic (green) or that co-expressed TH immunoreactivity (dark purple) and/or *Th* hybridization (light purple) throughout the anteroposterior extent of the zona incerta (ZI) in a *Th-cre;L10-Egfp* mouse (N = 3). Representative photomicrograph of the division between the medial and lateral ZI (yellow dashed line), which was defined by the lateral edge of the mammillothalamic tract (mtt) and fornix (fx; **c**). High magnification photomicrographs (from dashed outlined area in c) of *Egfp*-expressing cells that were ectopic (asterisk) or that co-expressed TH immunoreactivity (closed arrowheads; **1**) and/or *Th* hybridization (open arrowheads; **2**) within the medial (**d**) or lateral ZI (**e**). Proportion of *Egfp* cells in the medial (**1**) and lateral ZI (**2**) that expressed *Th* mRNA or TH immunoreactivity (**f**). Numeral inside the donut chart indicates the average number of cells counted within the ZI. Average cell counts and maps at each *Allen Reference Atlas* (*ARA*) level are based on at least two brains. Maps were produced using *ARA* templates (Dong, 2008) with reference to cytoarchitectural boundaries on Nissl-stained tissue. Panels with maps include the corresponding atlas level (bottom right), Bregma (*β*; top right), and brain region labels using formal nomenclature from the *ARA*. Scale bar: 50 μm (**c**); 20 μm (**d, e**). 3V, third ventricle; da, dopaminergic A13 group.

The vast majority (92 ± 4%) of *Egfp* ZI cells expressed *Th* mRNA, and only a few ZI cells (<1%) that expressed *Th* mRNA were not *Egfp*-labeled. Interestingly, only 27 ± 5% of *Th*-expressing *Egfp* cells also expressed TH immunoreactivity, and these TH-ir cells predominantly clustered within L67–69 only (**Figure 1a**, **b**). Furthermore, TH-ir, *Th*-, and *Egfp*-hybridized cells tended to be medially distributed within the ZI (**Figure 1b**). However, since we could not discern a Nissl-defined cytoarchitectural boundary within the ZI, we defined the medial and lateral ZI by a border along the lateral edge of the mammillothalamic tract and fornix (**Figure 1c**). The proportion of *Th*-hybridized *Egfp* cells were comparable between the medial (59 ± 5%; **Figure 1d1–d2**) and lateral ZI (41 ± 5%; p = 0.227; **Figure 1e1–e2**), but strikingly, the proportion of *Th*- and *Egfp*-hybridized cells that expressed TH immunoreactivity was significantly greater in the medial ZI (43 ± 8%; **Figure 1f1**) than lateral ZI (4 ± 0.6%; p = 0.038; **Figure 1f2**). Some ectopic *Egfp* cells (8 ± 3%) did not express either *Th* mRNA or TH immunoreactivity and were similarly scattered within the medial (3 ± 1%; **Figure 1f1**) and lateral ZI (14 ± 6%; p = 0.156; **Figure 1f2**).

### 3.2 Injection cases at medial ZI may contain low or high TH immunoreactivity

As *Th-cre* cells co-expressing TH-immunoreactivity were more prominent in the medial ZI, we delivered a Cre-dependent viral tracer encoding mCherry into the medial ZI of *Th-cre* or *Th-cre;L10-Egfp* mice (N = 7) in order to map the fiber projections from putative ZI DA cells, which were labeled by DsRed immunoreactivity. Importantly, when injected into *Th-cre;L10-Egfp* mice, all DsRed-ir cells colocalized with EGFP (**Figure 2a–b**) thus indicating the specificity of DsRed expression for *Th-cre* cells. To determine if TH-ir cells in the medial ZI contributed unique fiber projections, we selected one injection case each (i.e., Case 1 and 7) that labeled a high versus low proportion of TH-ir DsRed-labeled cells (**Table 4**). Overall, about 70% of virally transduced DsRed-ir cells were constrained to the ZI, but we excluded cases where less than 50 DsRed-ir cells were labeled per brain series (i.e., Case 4, 5, and 6). We prioritized injection cases with a higher number of *Th-cre* cells transduced, thus we mapped and compared Case 1 and Case 7, which reflected cases with the lowest (17%) and highest (53%) percentage of TH-ir DsRed-labeled cells, respectively (**Table 4**).

**Figure 2.**
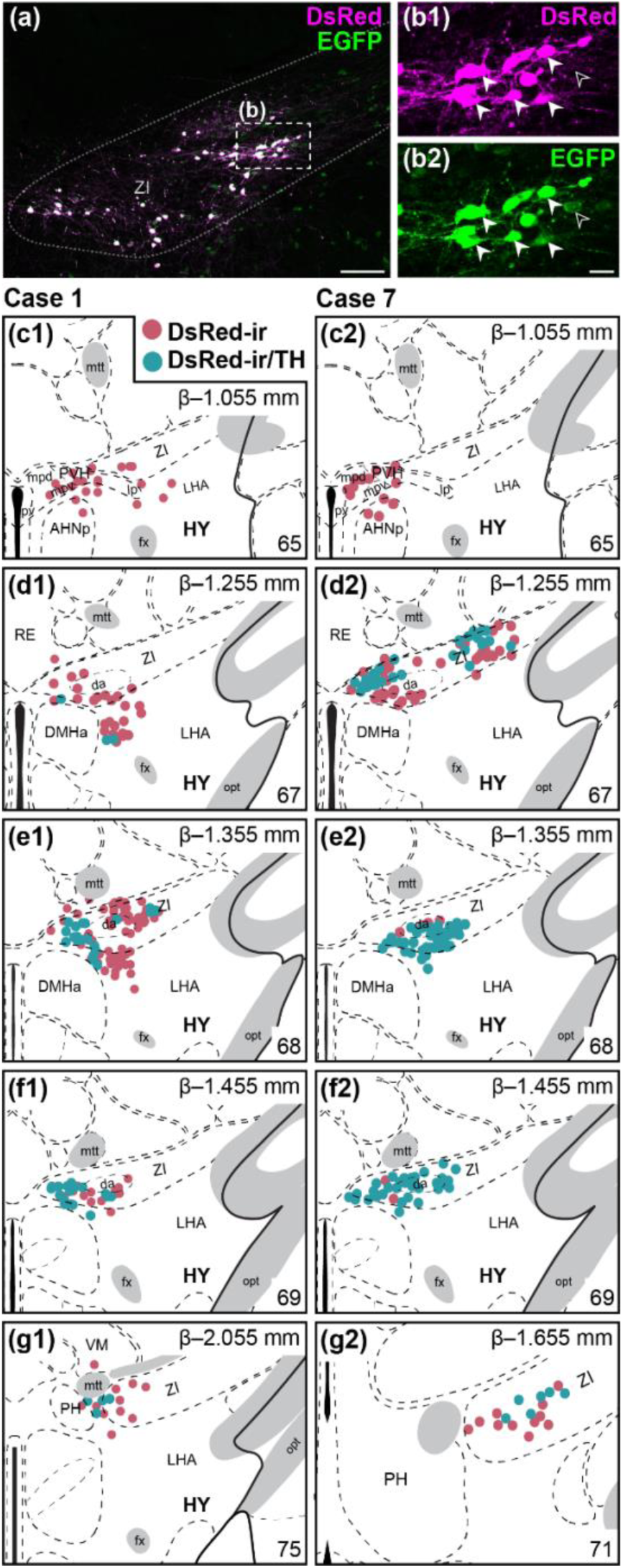
Virally-transduced DsRed-immunoreactive EGFP cells co-expressed TH primarily in the medial ZI. Representative photomicrograph of a cre-dependent virus encoding DsRed delivered into the ZI of a *Th-cre;L10-Egfp* mouse (**a**). High magnification photomicrograph (from dash outlined area in **a**) demonstrating EGFP cells (open arrowhead) co-expressing DsRed immunoreactivity (white arrowhead) in the ZI (**b**). Mapped distribution of DsRed-immunoreactive cells (red circles) co-expressing TH (blue circles) in injection Case 1 (**c1–g1**) and Case 7 (**c2–g2**). Cells were mapped onto *Allen Reference Atlas* (*ARA*) templates (Dong, 2008) with reference to cytoarchitectural boundaries on Nissl-stained tissue. Panels with maps include the corresponding atlas level (bottom right), Bregma (*β*; top right), and brain region labels using formal nomenclature from the *ARA*. Scale bar: 50 μm (**a**); 20 μm (**b**). ZI, zona incerta.

**Table 4.** List of injection cases at ZI *Th-cre* cells.

| Case<sup>a</sup> | ARA Range<sup>b</sup> | DsRed-ir cells<sup>c</sup> |  | TH-ir colocalization<sup>e</sup> |
| --- | --- | --- | --- | --- |
|  |  | Total | ZI<sup>d</sup> |  |
| 1 | 65–71 | 132 | 68% | 17% |
| 2 | 63–67 | 69 | 95% | 50% |
| 3 | 63–69 | 76 | 70% | 47% |
| 4 | 63–69 | 18 | 100% | 33% |
| 5 | 62–70 | 35 | 85% | 51% |
| 6 | 62–75 | 41 | 89% | 37% |
| 7 | 65–75 | 86 | 85% | 53% |

<sup>a</sup> Injections in *Th-cre* or *Th-cre;L10-Egfp* mice designated T and TG, respectively.
<sup>b</sup> Atlas levels defined by *Allen Reference Atlas* templates (Dong, 2008).
<sup>c</sup> Virally transduced DsRed-immunoreactive (-ir) cells counted unilaterally from one of four brain series sliced from the whole brain.
<sup>d</sup> Percent of DsRed-ir cells constrained to the ZI.
<sup>e</sup> Percent of DsRed-ir cells that coexpress tyrosine hydroxylase (TH) immunoreactivity.

### 3.3 Mapping intersecting points to define common spatial targets of ZI *Th-cre* cells

We mapped DsRed-labelled fibers from Case 1 and Case 7 onto *ARA* brain atlas templates (Dong, 2008) and recorded the amount of fiber coverage, which we depicted in pixels, across the major divisions of the brain (Wang et al., 2020a; Allen Institute for Brain Science, 2011). The fiber coverage within each brain division was strongly correlated between Case 1 and Case 7 (r = 0.994; p < 0.0001; **Figure 3a**), thus we overlaid the mapped fiber distribution from Case 1 and Case 7 to determine their points of intersection to define common spatial targets of medial ZI *Th-cre* cells.

**Figure 3.**
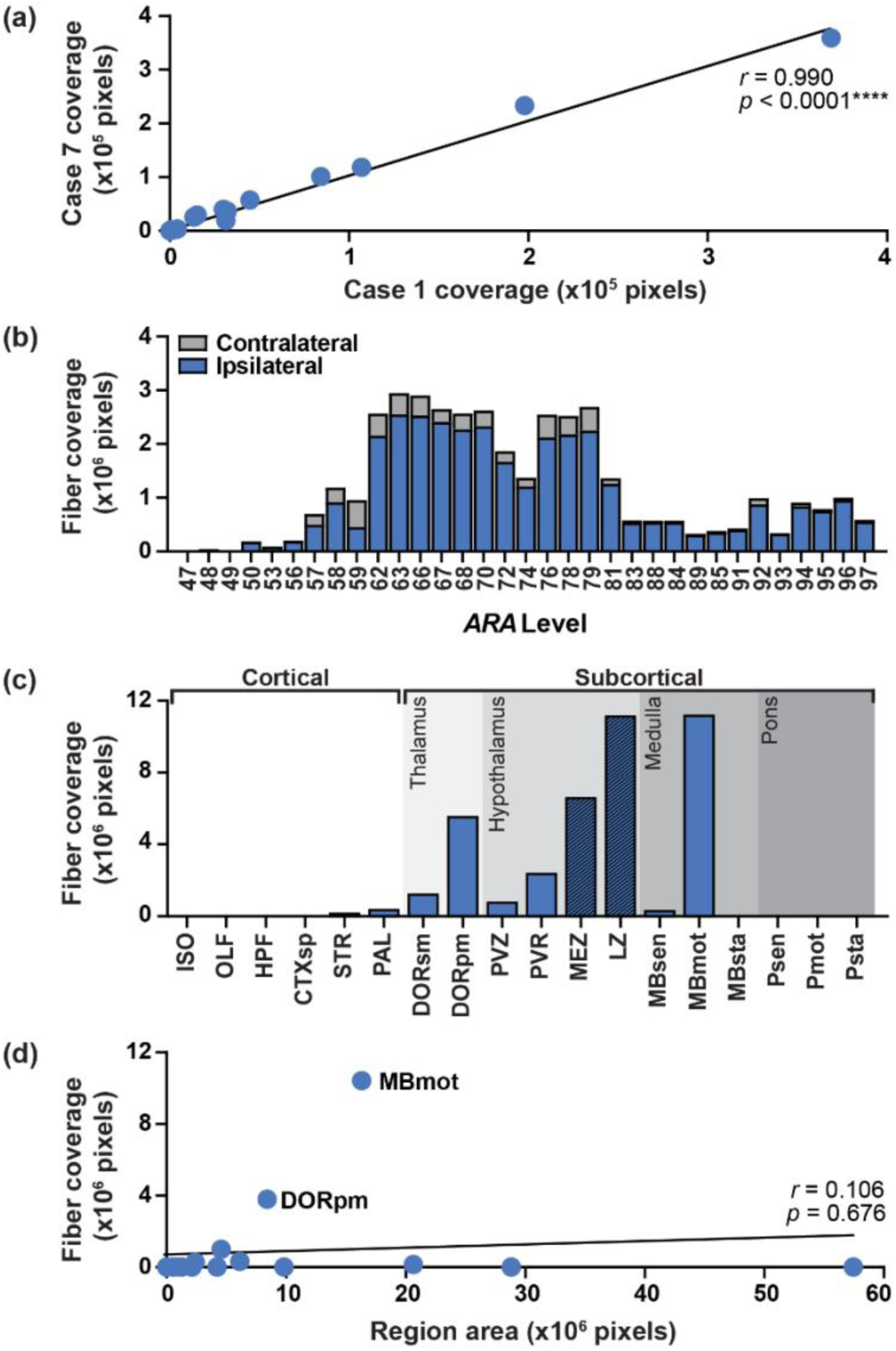
Motor-related midbrain comprised the most abundant fiber projections from *Th-cre* ZI cells. Correlation of fiber coverage (in pixels) between injection Case 1 and Case 7 across major brain divisions (see **Supplemental Figure 1**) as defined by the Allen Mouse Brain Atlas (Wang et al., 2020a; Allen Institute for Brain Science, 2011; **a**). The amount of contralateral and ipsilateral fiber coverage formed by intersecting points between Case 1 and Case 7 throughout the entire brain (**b**). Amount of fiber coverage from intersecting points across each major division of the brain (**c**) with hatched bars indicating regions comprising the injection site. Correlation of fiber coverage from intersecting points and total region area (in pixels; **d**). Correlational analyses were determined by Pearson’s *r*, where ****, p < 0.0001. CTXsp, cortical subplate; DORpm, polymodal-association cortex related regions of the thalamus; DORsm, sensory-motor cortex related regions of the thalamus; HPF, hippocampal formation; ISO, isocrotex; LZ, hypothalamic lateral zone; MBmot, motor related regions of the midbrain; MBsen, sensory related regions of the midbrain; MBsta, behaviour state related regions of the midbrain; MEZ, hypothalamic medial zone; OLF, olfactory regions of the cortex; PAL, pallidum; Pmot, motor related regions of the pons; Psen, sensory related regions of the pons; Psta, behaviour state related regions of the pons; PVR, periventricular region; PVZ, periventricular zone; STR, striatum.

Our intersecting analysis focused on *ARA* levels that spanned between +0.750 mm and −4.380 mm from begma (L44–97) and on the hemisphere ipsilateral to the injection site where most DsRed-ir fibers were distributed (**Figure 3b**). Rostrally, DsRed-ir fibers appeared anterior to the genu of the corpus callosum (L47, +0.750 mm from bregma) and extended caudally into the posterior hindbrain. Unfortunately, sections caudal to L97 were not collected for Case 1, thus DsRed-labelled fibers posterior to L97 were mapped from Case 7 only.

DsRed-ir fibers were largely restricted to subcortical structures in the thalamus, hypothalamus, and medulla (**Figure 3c**). Expectedly, we found strong DsRed labeling in the hypothalamic medial (MEZ) and lateral zone (LZ) where the injection site was localized. The polymodal association cortex-related thalamus (DORpm) and motor-related midbrain (MBmot) contained the greatest fiber coverage among major brain divisions away from the injection site (**Figure 3c**). We considered whether the abundance of DsRed-labelled fibers in the DORpm and MBmot was related to the size of these brain divisions, but we found that the prevalence of fibers in a region was not related to the brain area (r = 0.11; p = 0.676; **Figure 3d**). Large brain divisions like the isocortex and striatum comprised only few DsRed-ir fibers, thus smaller brain divisions like the DORpm and MBmot that comprised high intersecting fiber coverage were especially notable (**Figure 3d**).

### 3.4 Mapping brainwide *Th-cre* fiber projections

The distribution patterns of DsRed-ir fibers were highly comparable between cases across all major brain divisions and most individual brain regions (**Figure 4**). We used a density scale (+++, high; ++, moderate; +, low; −, very low or none) to report semi-qualitative estimates of fiber distribution in each brain region (**Figure 5**), and we provided a comprehensive summary of the projection patterns across all major brain divisions using the hierarchical structure (Wang et al., 2020a) and nomenclature (Allen Institute for Brain Science, 2011), which was set forth by the *Allen Reference Atlas*, for each injection case in **Table 5**. Where possible, we made note of terminal boutons, which appeared as puncta clusters around a soma, to distinguish them from fibers-of-passage along the dorsoventral axis, which presented as puncta arranged in a continuous line (**Figure 6**). We provided side-by-side comparisons of the distribution of DsRed-ir fibers from Case 1 (**Figure 7a1–ff1**), Case 7 (**Figure 7a2–ff2**), and their points of intersection (**Figure 7a3–ff3**) across all levels available in our dataset.

**Figure 4.**
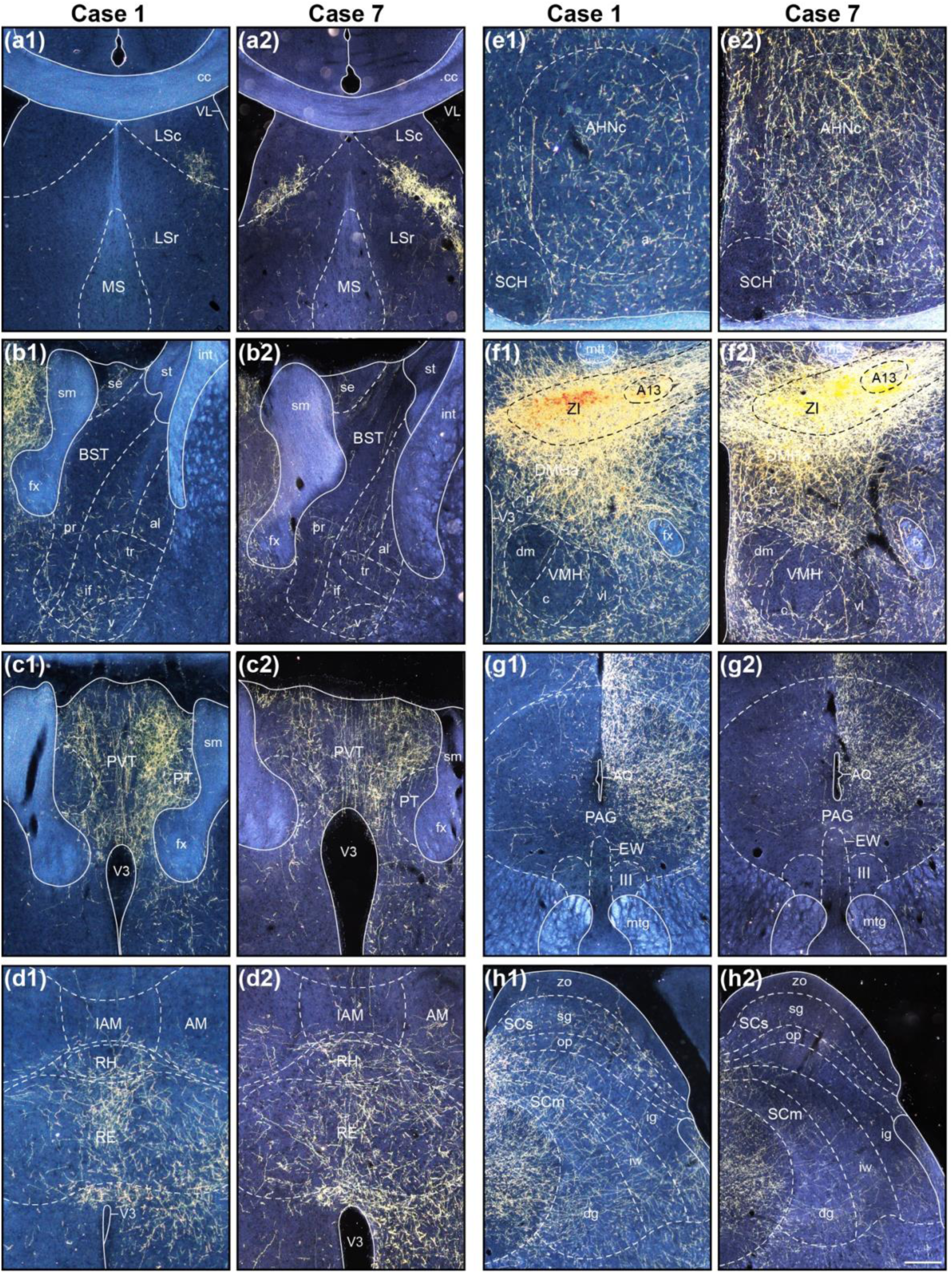
DsRed-labeled fibers from *Th-cre* ZI cells are abundant throughout the brain. Representative darkfield photomicrographs of DsRed-immunoreactive fibers from injection Case 1 (**a1–h1**) and Case 7 (**a2–h2**). Scale bar, 50 μm. A13, dopaminergic A13 group; AHN, anterior hypothalamic nucleus; AHNa, anterior part; AHNc, central part; AM, anteromedial nucleus of the thalamus; AQ, cerebral aqueduct; BST, bed nucleus of the stria terminalis; BSTal, BST anterolateral area; BSTif, BST interfasicular nucleus; BSTpr, BST principle nucleus; BSTse, BST strial extension; BSTv, ventral nucleus; cc, corups callosum; DMH, dorsomedial hypothalamic nucleus; DMHa, DMH anterior part; DMHp, DMH posterior part; EW, Edinger-westphal nucleus; fx, fornix; IAM, interanteromedial nucleus of the thalamus; III, oculomotor nucleus; int, internal capsule; LS, lateral septal nucleus; LSc, LS caudal (caudodorsal) part; LSr, LS rostral (rostroventral) part; MS, medial septal nucleus; mtg, mammillotegmental tract; mtt, mammillothalamic tract; PAG, periaqueductal grey area; PT, paratenial nucleus; PVT, paraventricular nucleus of the thalamus; RE, nucleus of reuniens; RH, rhomboid nucleus; SCH, suprachiasmatic nucleus; SCm, superior colliculus, motor-related; SCdg, SCm deep gray layer; SCig, SCm intermediate gray layer; SCiw, SCm intermediate white layer; SCs, superior colliculus, sensory related; SCop, SCs optic layer; SCsg, SCssuperficial gray layer; SCzo, SCs zonal layer; sm, stria medullaris; st, stria terminalis; tr, transverse nucleus; V3, third ventricle; VL, lateral ventricle; VMH, ventromedial hypothalamic nucleus; VMHc, central part; VMHdm, dorsomedial part; VMHvl, ventrolateral part; ZI, zona incerta.

**Figure 5.**
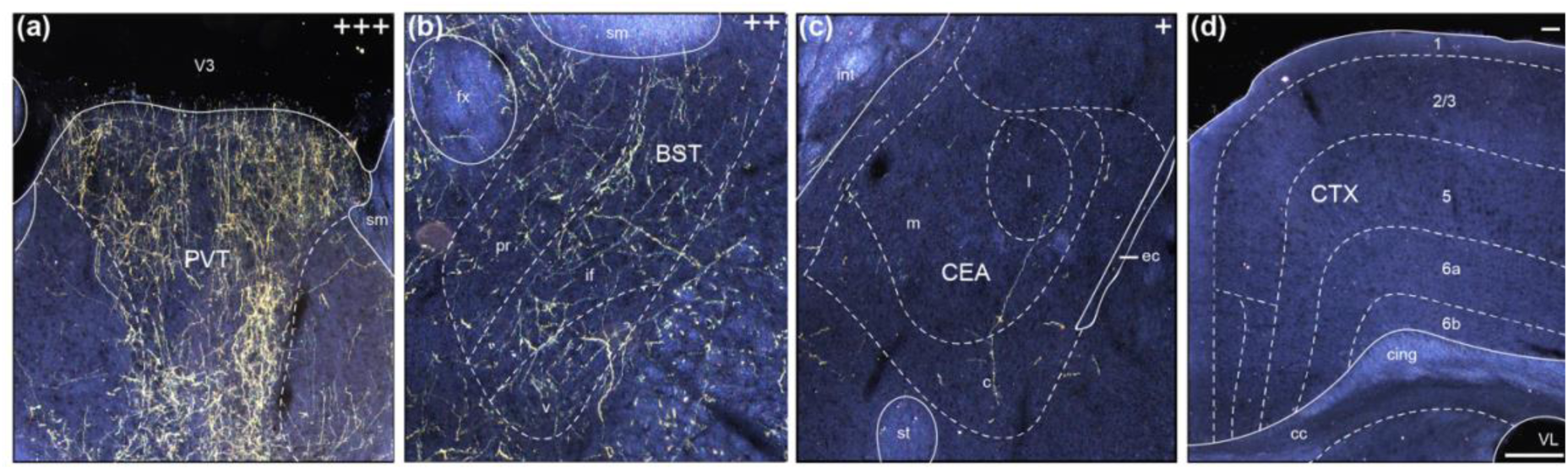
Difference in density of DsRed-labeled fibers from *Th-cre* ZI cells. Representative darkfield photomicrographs of DsRed-immunoreactive fibers providing high (+++; **a**), moderate (++; **b**), low (+; **c**), and very little or no fiber coverage (−; **d**) following cre-mediated viral transduction of *Th-cre* cells in the medial ZI. Scale bar, 50 μm. 1, layer 1; 2/3, layer 2/3; 5, layer 5; 6a, layer 6a; 6b, layer 6b; sm, stria medullaris; st, stria terminalis; BST, bed nucleus of the stria terminalis; BSTif, BST interfasicular nucleus ; BSTpr, BST principle nucleus; BSTv, BST ventral nucleus; cc, corups callosum; CEA, central amygdala; CEAl, CEA lateral part; CEAm, CEA medial part; CEAc, CEA central part; cing, cingulum bundle; CTX, cortex; ec, external capsule; fx, fornix; int, internal capsule; PVT, paraventricular nucleus of the thalamus; V3, third ventricle; VL, lateral ventricle.

**Figure 6.**
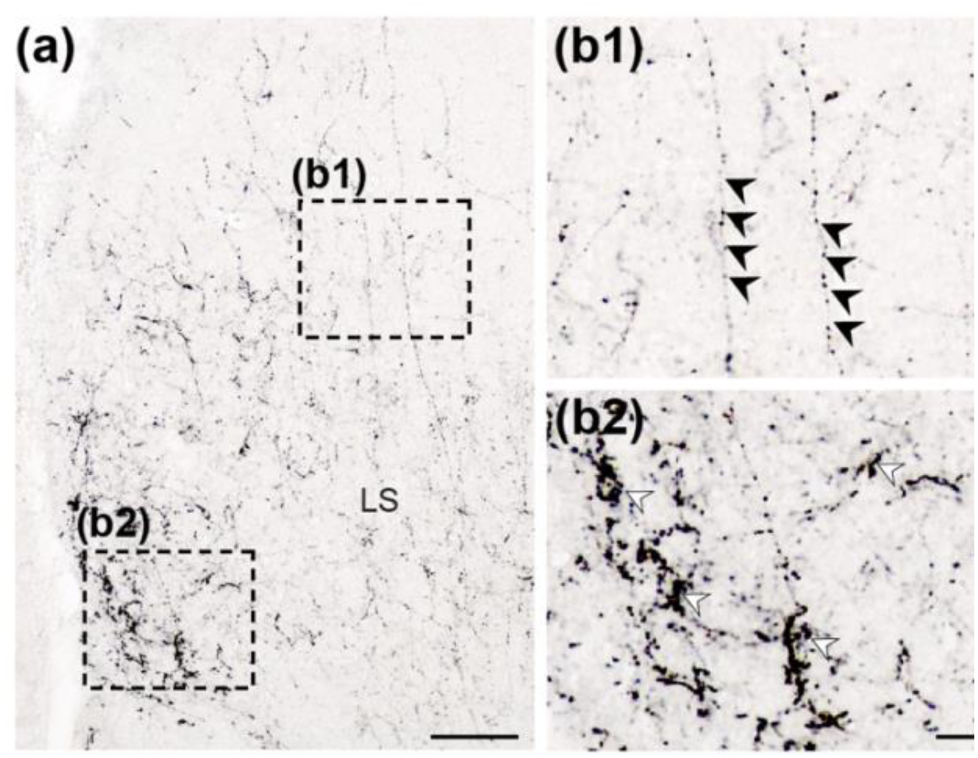
Distinguishing between fibers-of-passage and terminal buttons from *Th-cre* ZI cells. Representative brightfield photomicrographs of DsRed-immunoreactive fibers from cre-mediated *Th-cre* ZI cells (**a**) revealing puncta arranged in a continuous path like fibers-of-passage (**b1**, black arrow heads) or that formed clusters around a soma (**b2**, white arrowheads). Scale bar: 50 μm (**a**); 20 μm (**b**). LS, lateral septum.

**Figure 7.**
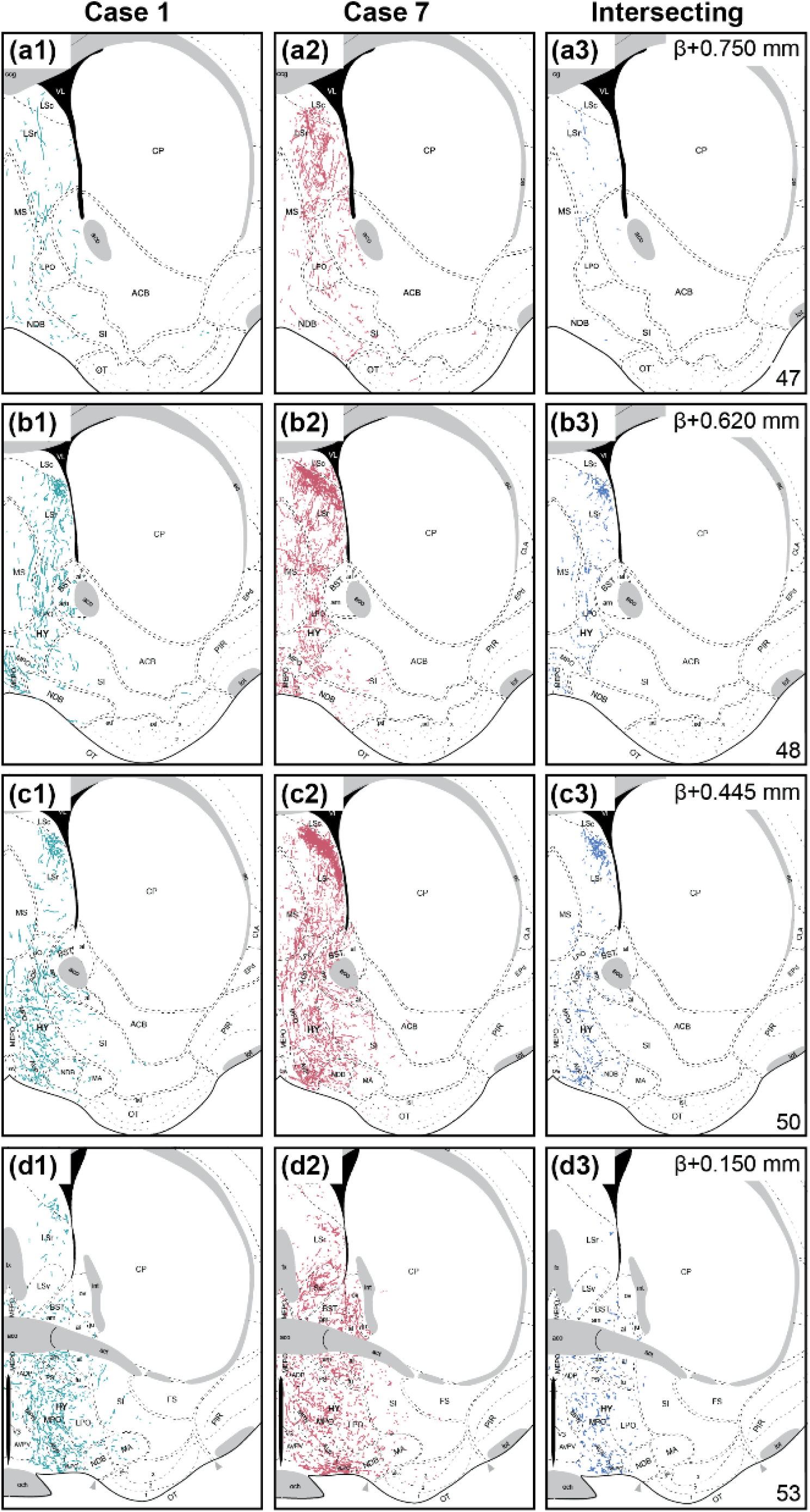
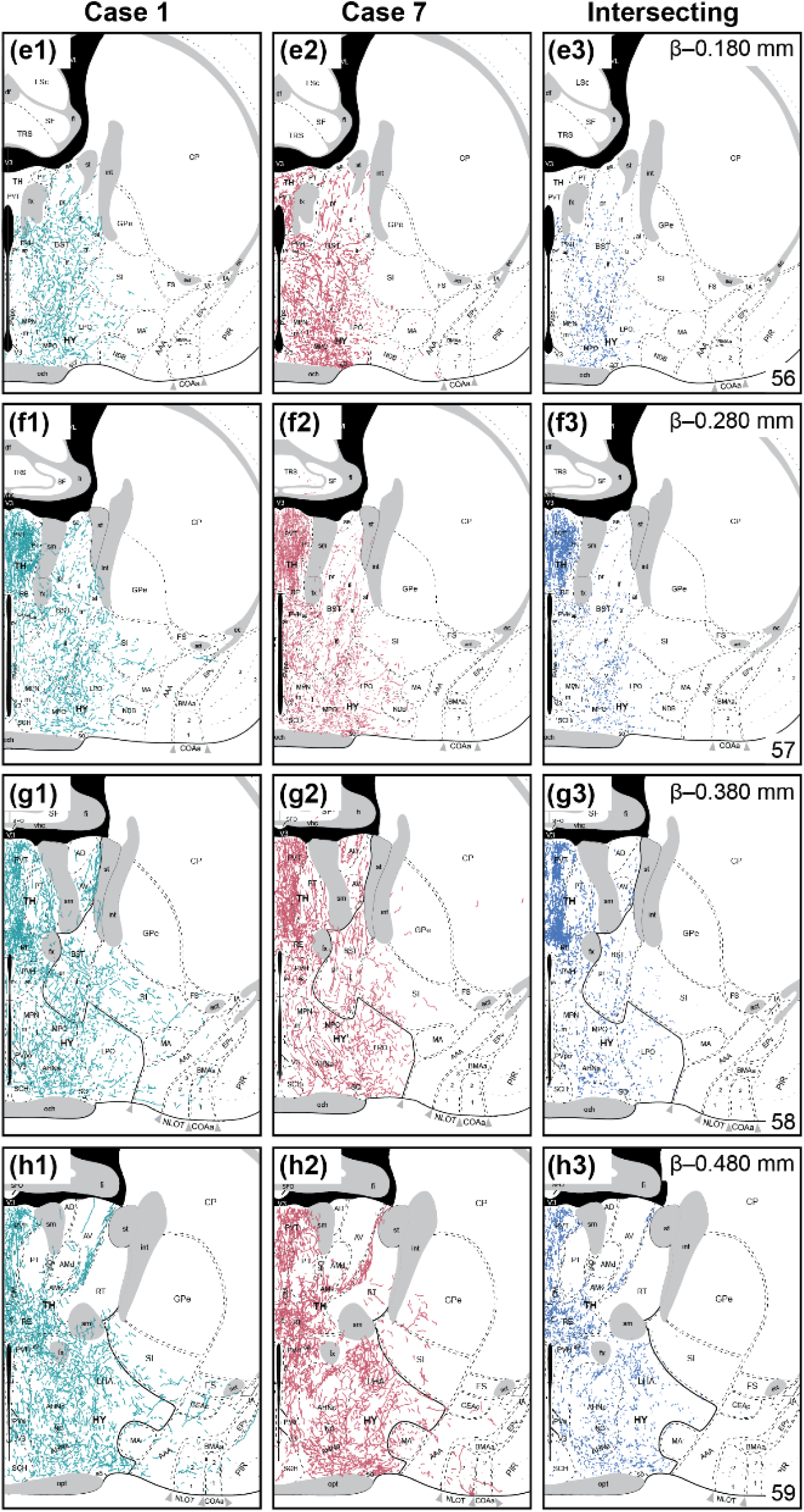
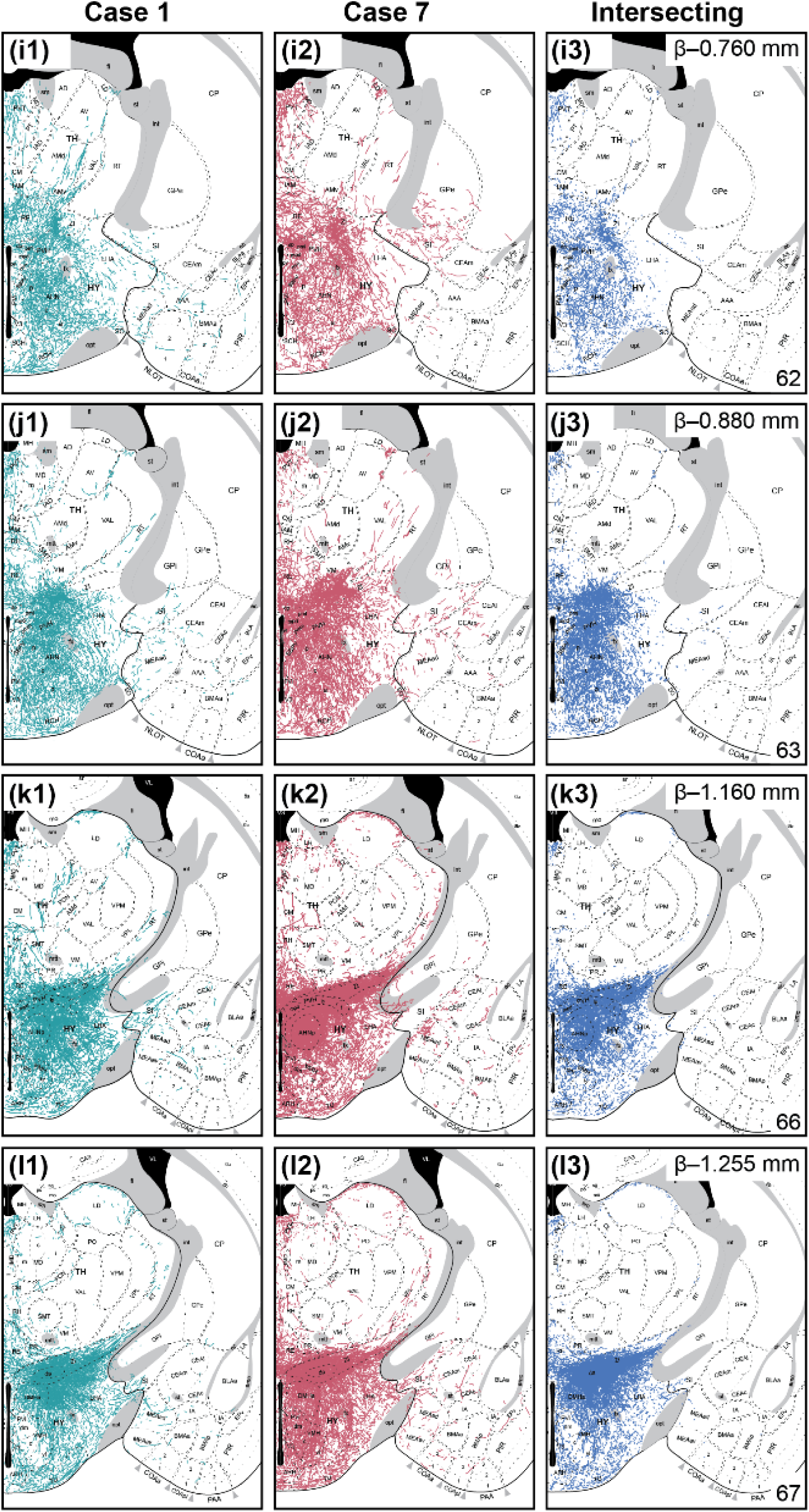
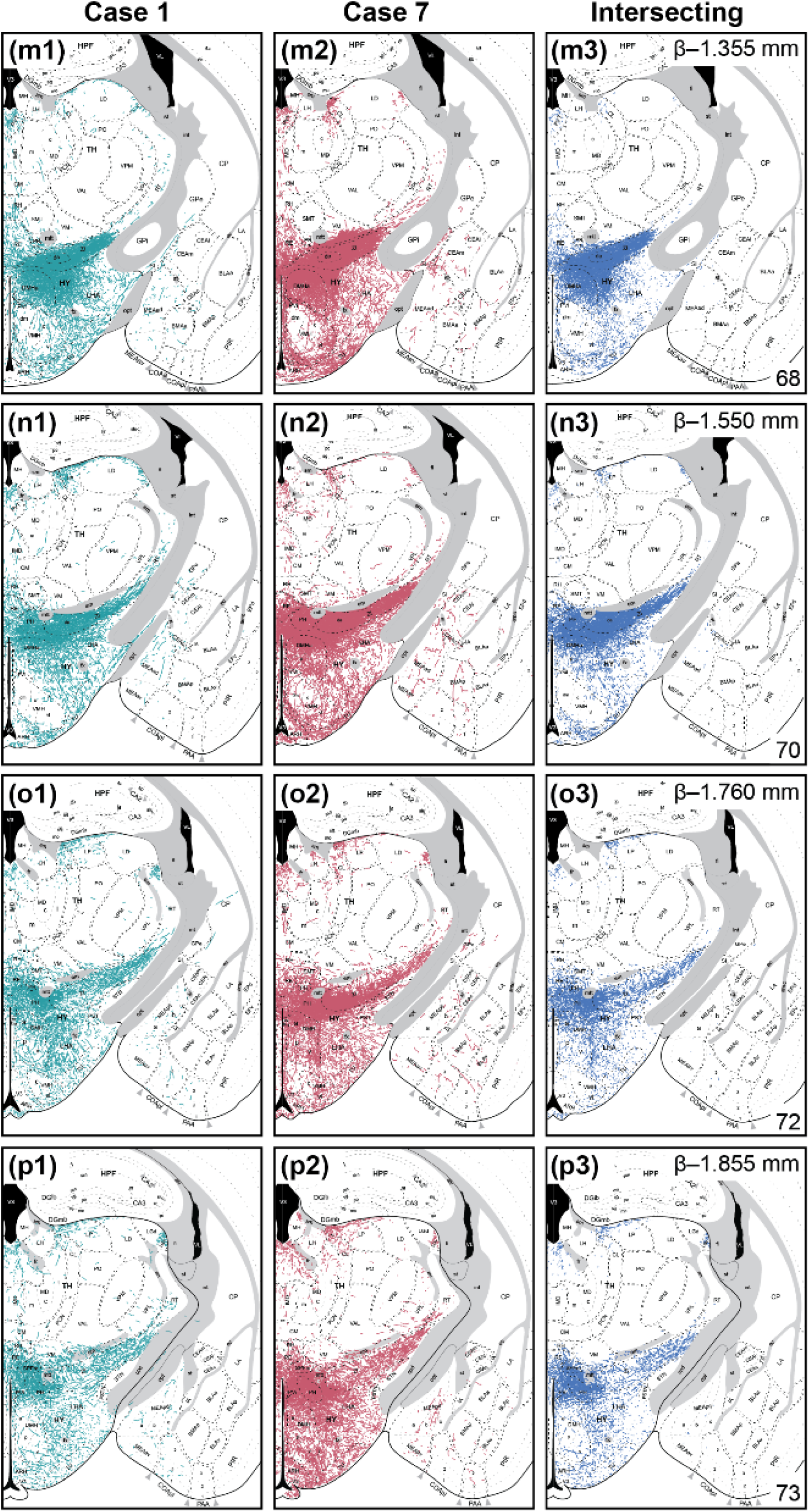
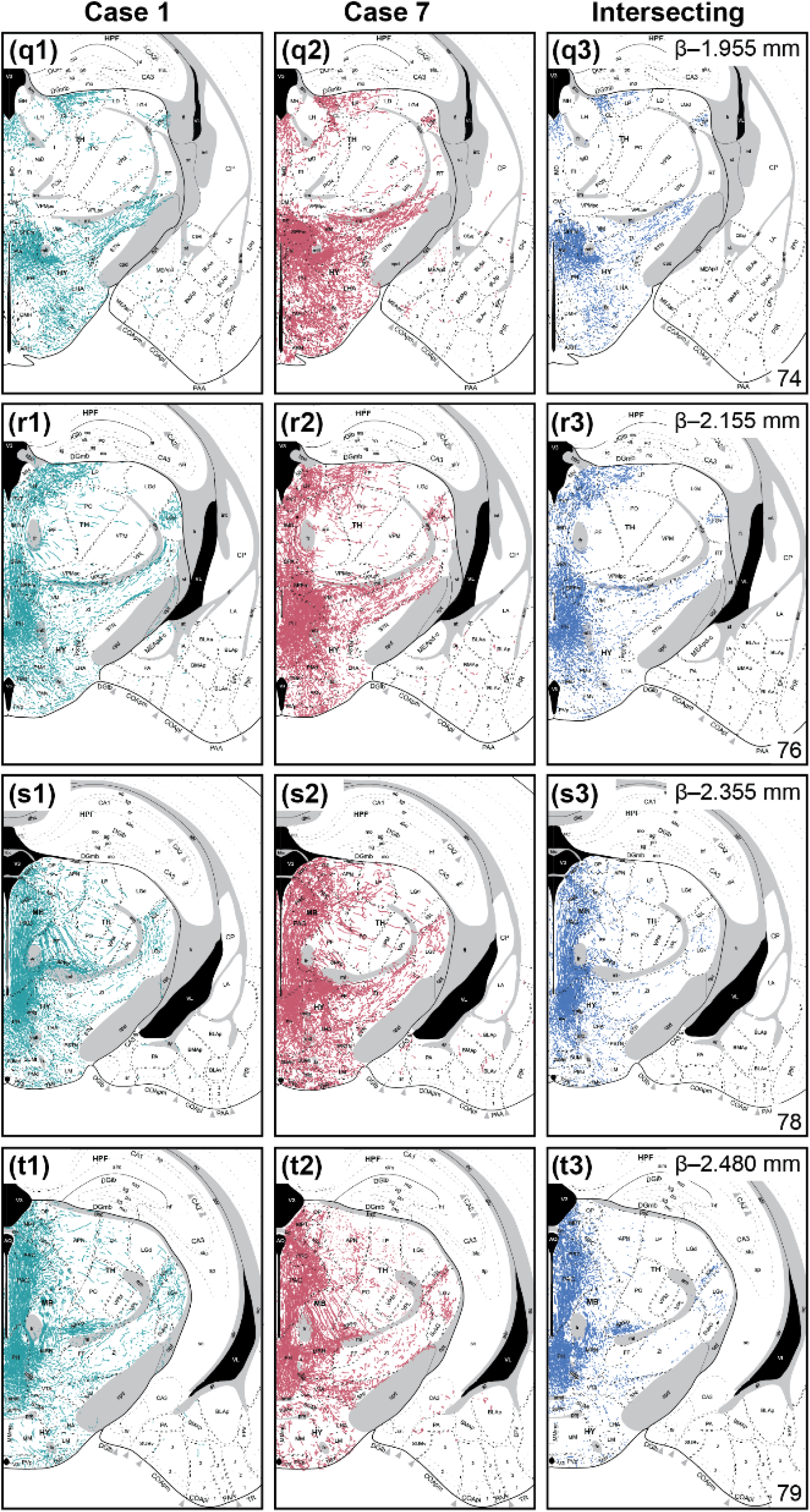
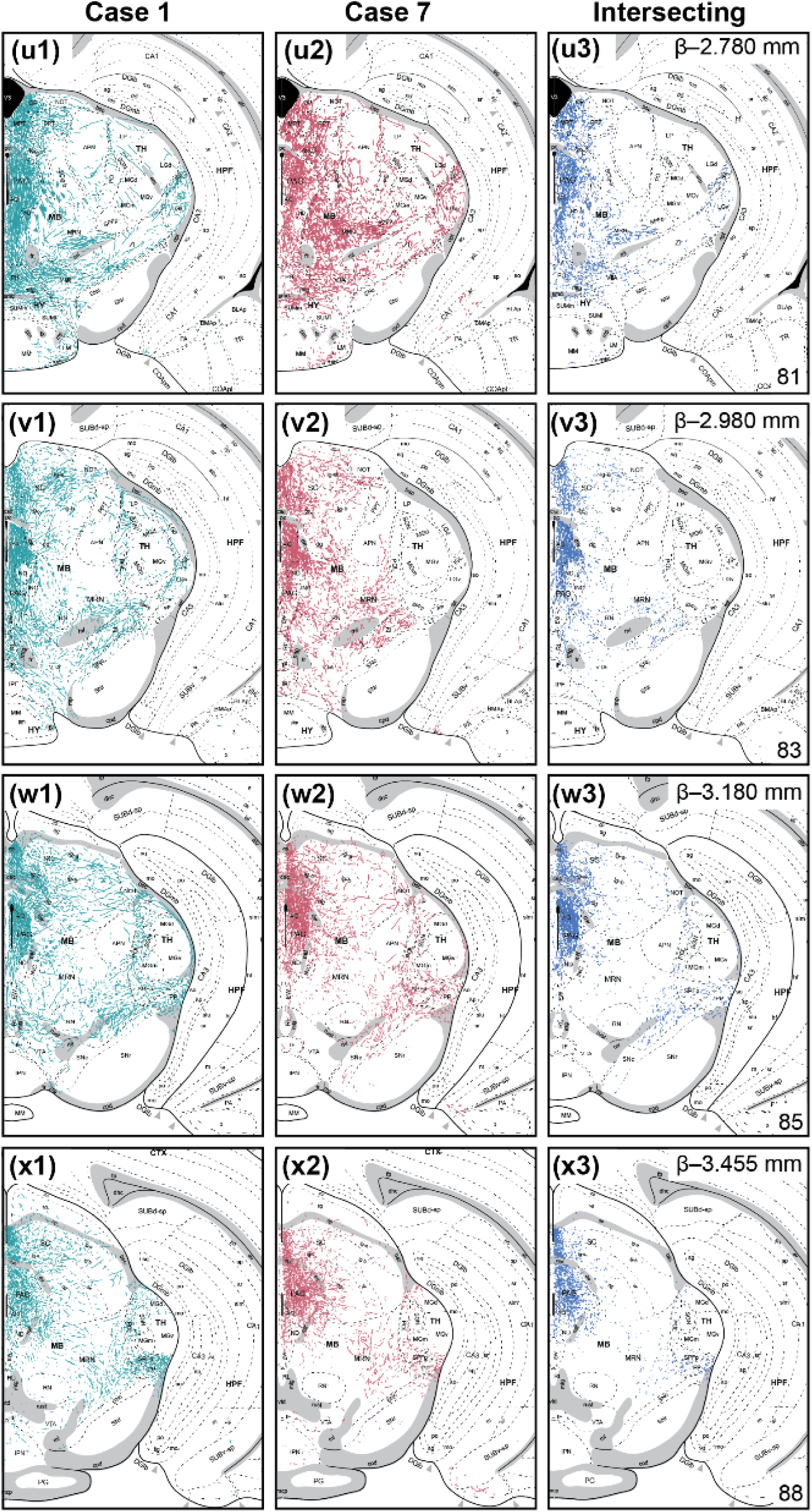
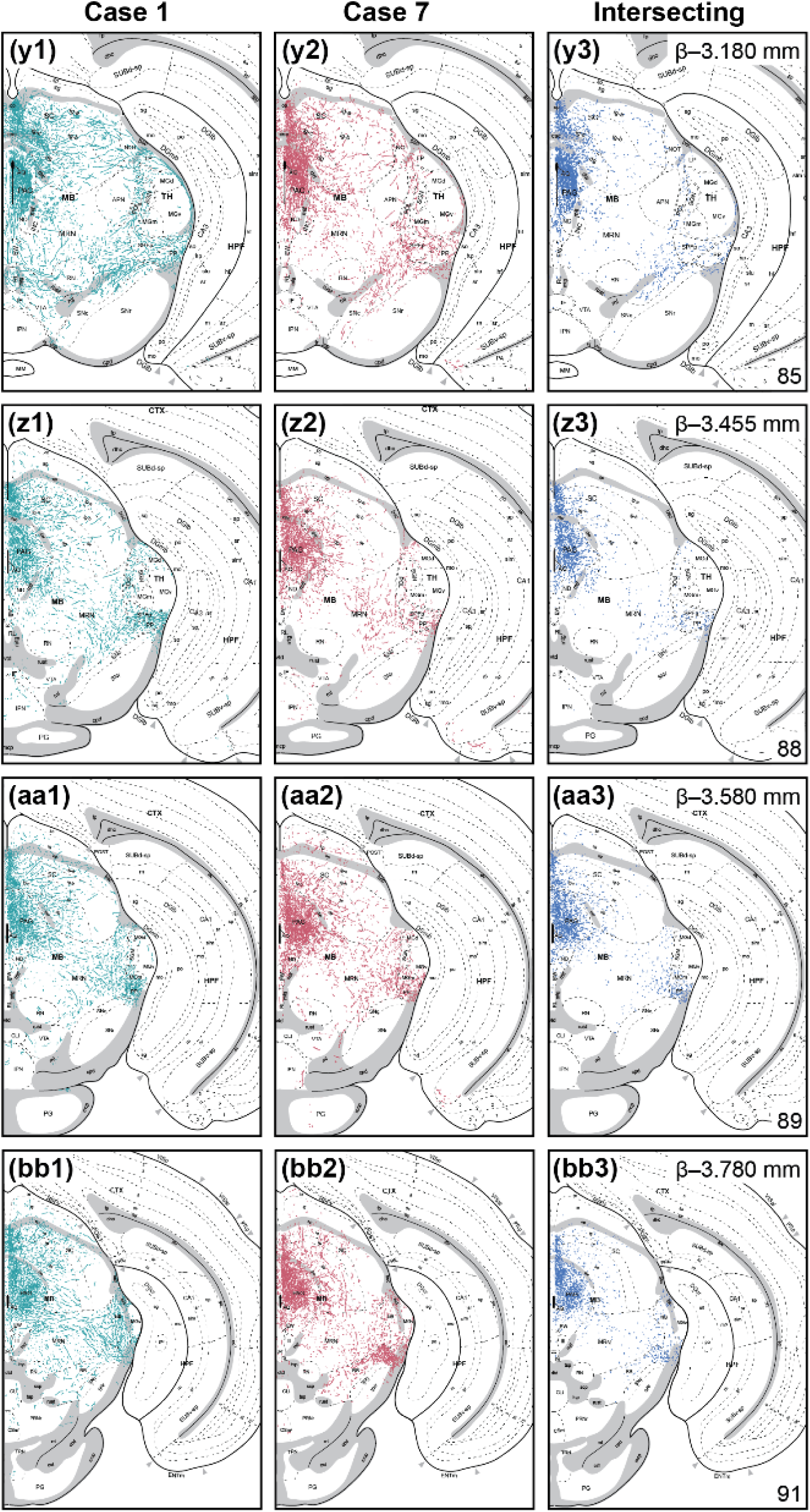
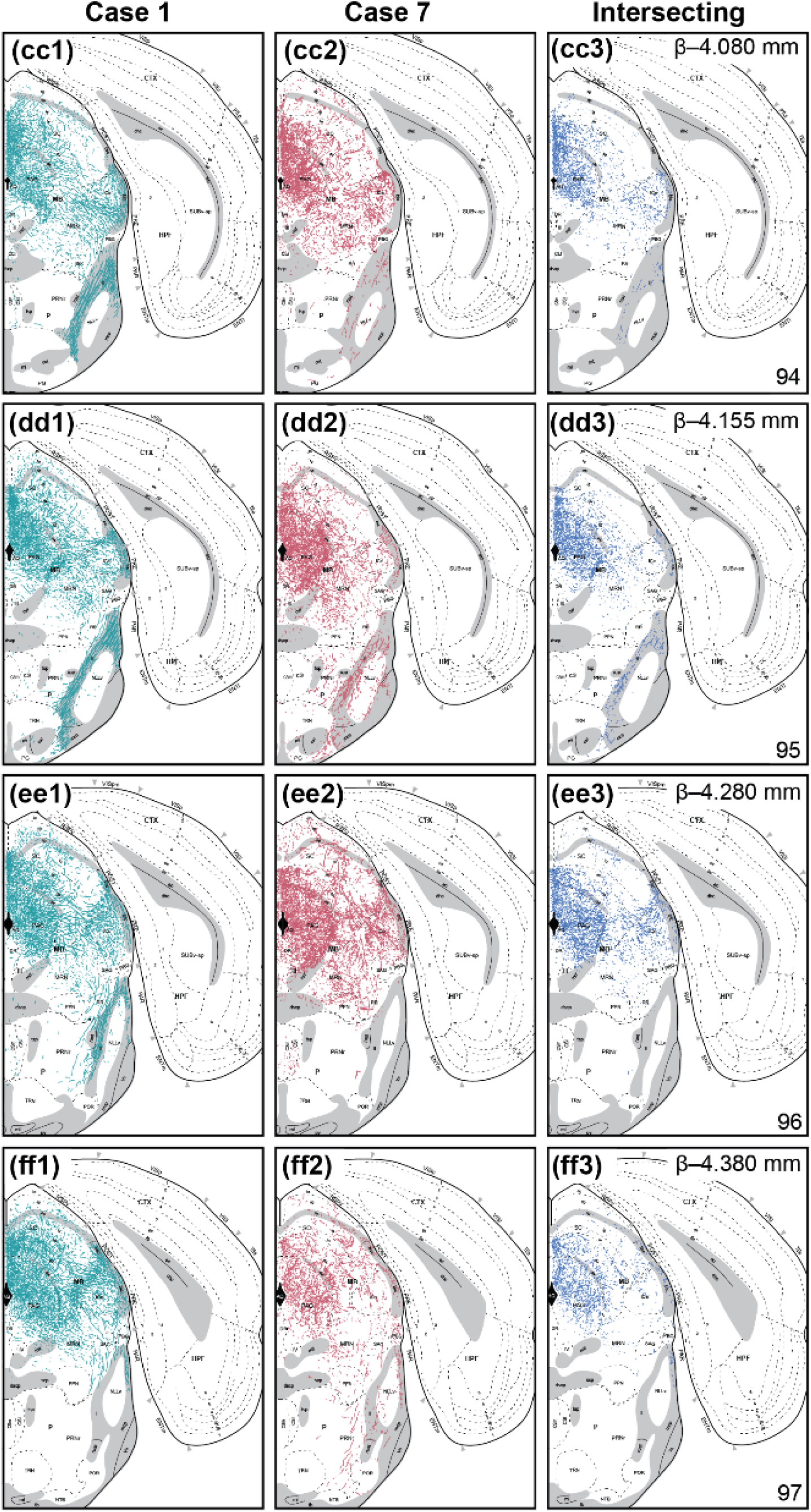
Distribution of fiber projections from *Th-cre* ZI cells from the forebrain to midbrain. Mapped distribution of fibers from injection Case 1 (**a1–ff1**), Case 7 (**a2–ff2**), and the points of intersection between both injection cases (**a3–ff3**). Traced fibers were overlaid onto *Allen Reference Atlas* (*ARA*) templates (Dong, 2008) with reference to cytoarchitectural boundaries on Nissl-stained tissue. Panels include the corresponding atlas level (bottom right), Bregma (*β*; top right), and brain region labels using formal nomenclature from the *ARA*.

**Table 5.**
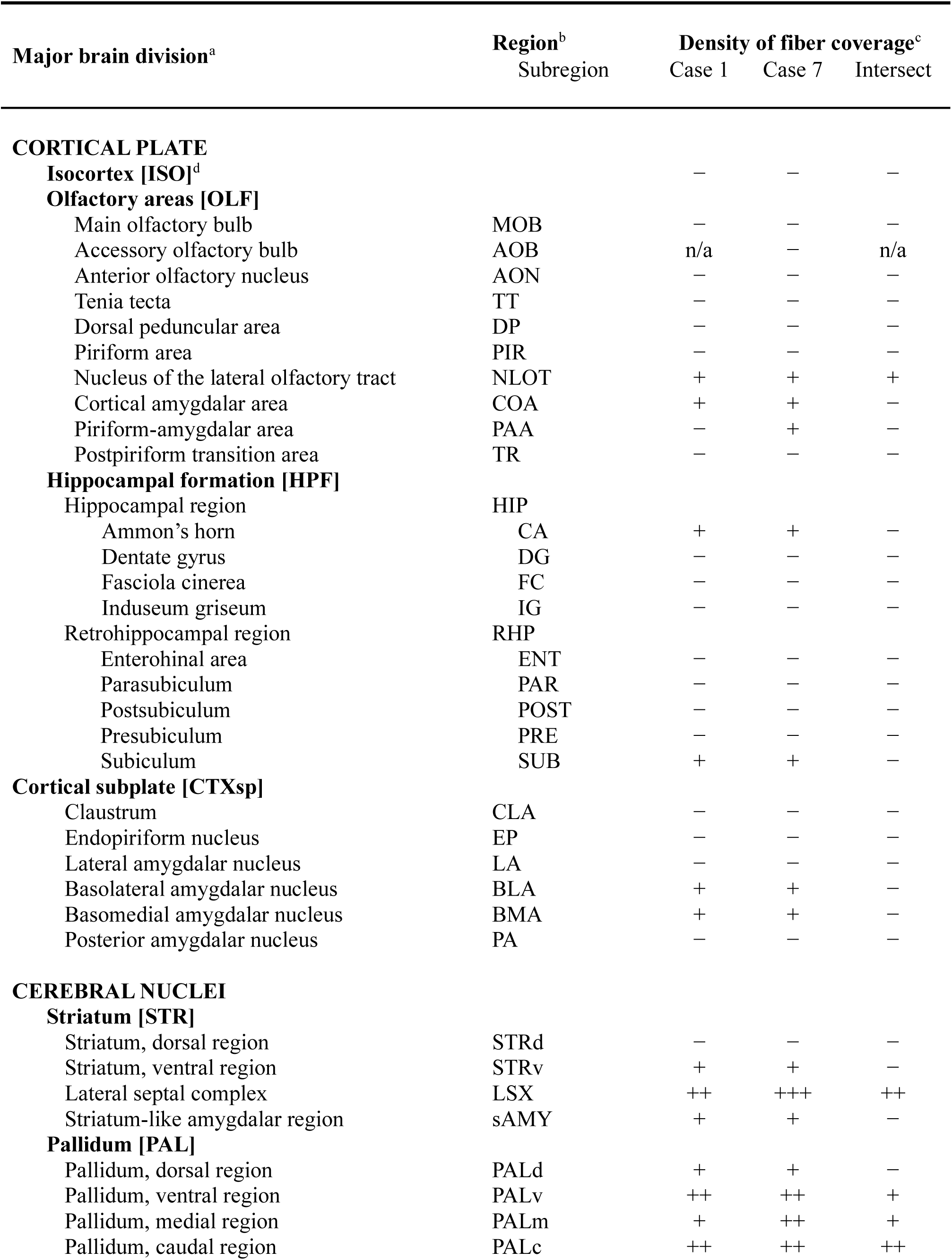
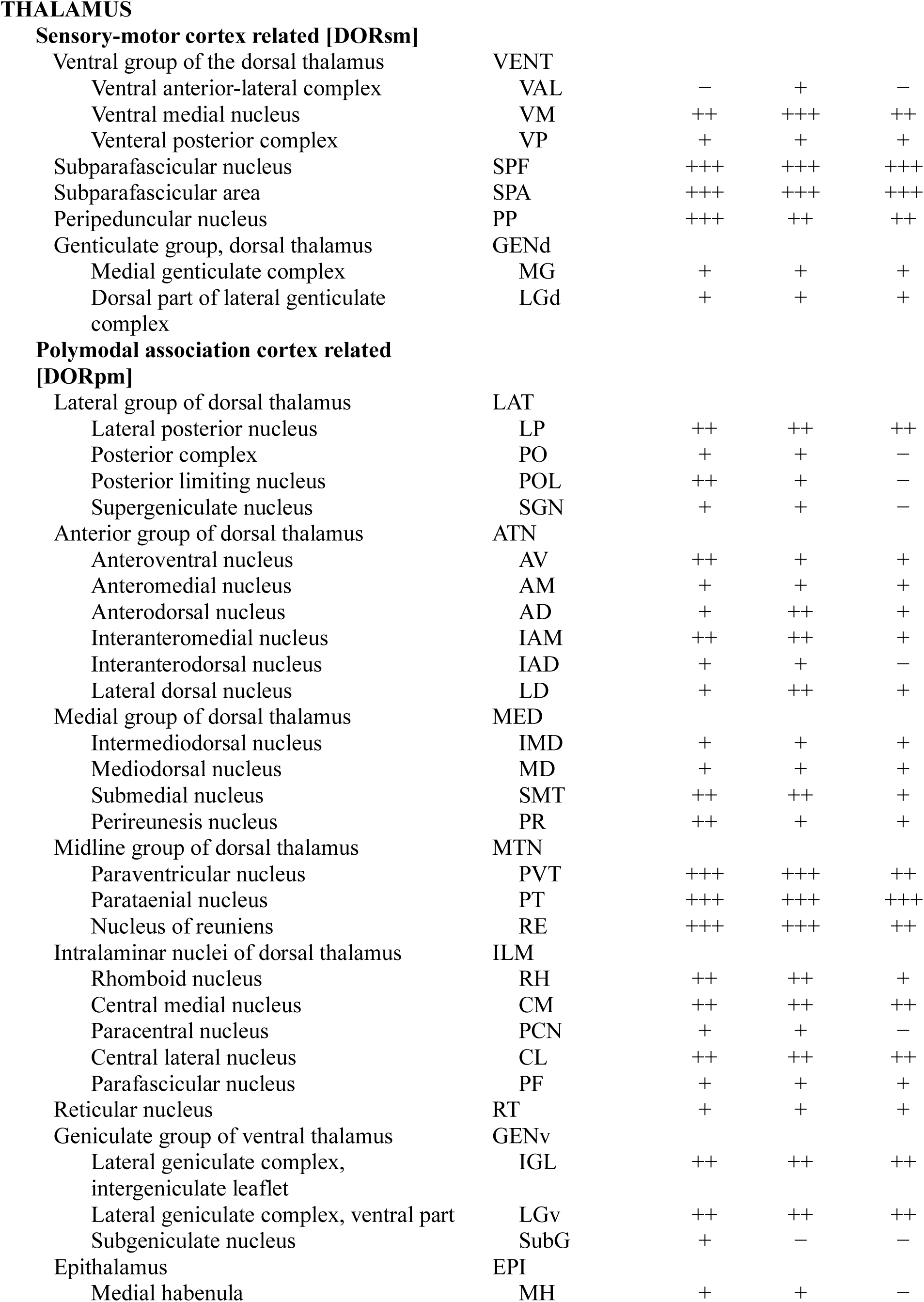
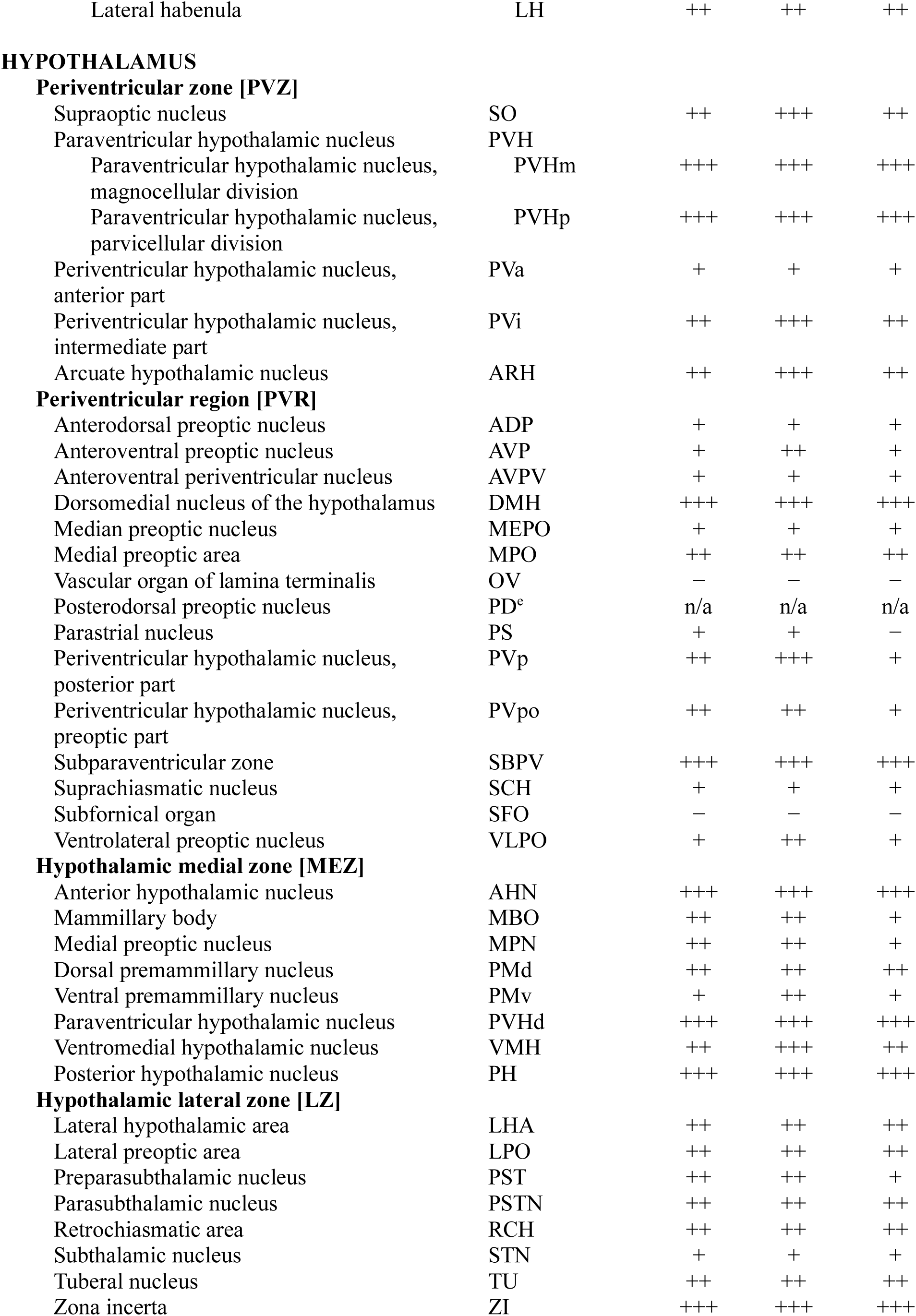
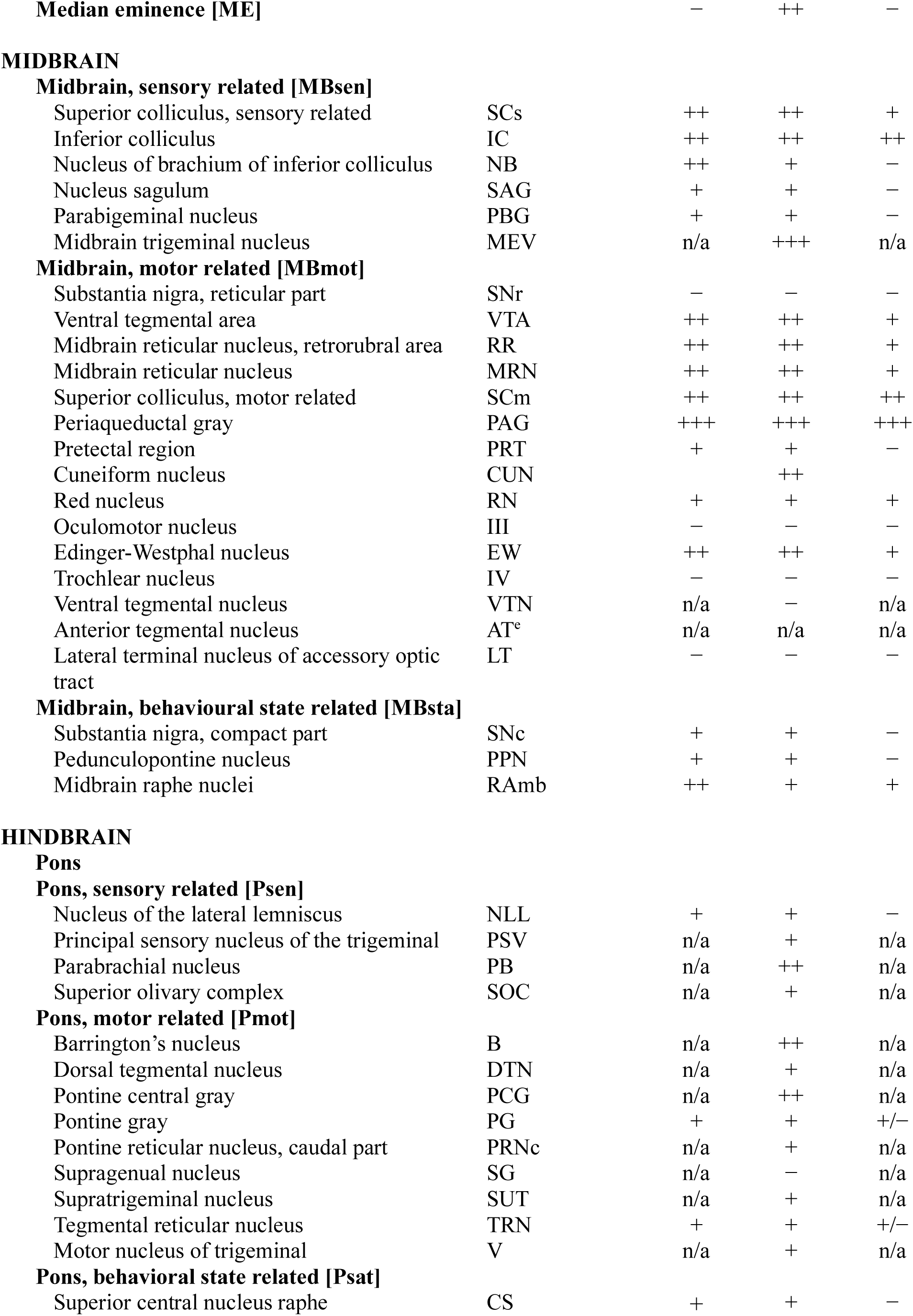
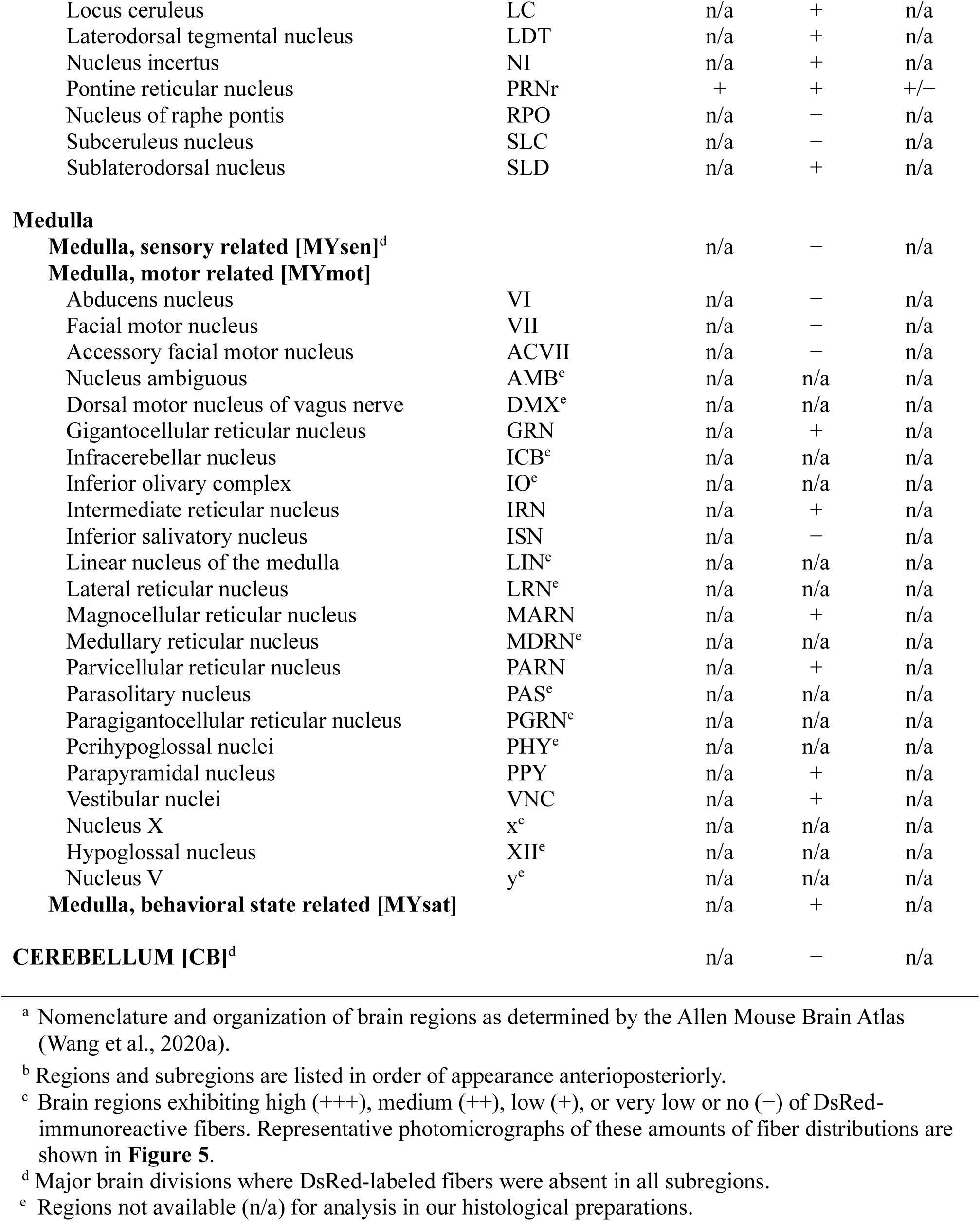
Qualitative distribution of fiber projections from ZI *Th-cre* cells.

#### 3.4.1 Cerebral cortex

DsRed-labelled fibers within the cerebral cortex were spare relative to all other major brain divisions analyzed (**Table 5**).

##### Cortical plate (CTXpl)

Subregions within the CTXpl including the isocortex, olfactory areas, and hippocampal formation contained few fibers. Although limited, fibers were observed in regions comprising olfactory areas including the lateral olfactory tract (NLOT; **Figure 7g–j**), cortical amygdalar area (COA; **Figure 6e–w**), and piriform-amygdalar area (PAA; **Figure 7k–t**). However, this may not be reflected in the intersecting maps due to their sparsity (**Table 5**). The remaining olfactory areas were completely devoid of fibers including the piriform area (PIR; **Figure 7a–u**), piriform-amygdalar area (PAA; **Figure 7k–t**), postpiriform transition area (TR; **Figure 7t–y**), main olfactory bulb (MOB), anterior olfactory nucleus (AON), tenia tecta (TT), and dorsal peduncular area (DP) (**Table 5**). Case 1 also revealed no fibers in the accessory bulb (AOB; **Table 5**), however we were unable to capture the AOB in Case 7 and subsequently, the intersecting maps.

In the hippocampal formation and retrohippocampal formation, DsRed-labelled fibers were only present in Case 7 posteriorly (L78–91; **Figure 7s2–z2** and appeared to terminate within the ventral portion of the Cornu Ammonis (CA) 1 (**Figure 7u2–v2**), CA3 (**Figure 7t2**), and ventral subiculum (SUBv; **Figure 7v2–y2**). The isocortex lacked DsRed-labelled fibers in both Case 1 and Case 7 (**Table 5**).

#### Cortical subplate (CTXsp)

There were few overlapping fibers along the anterioposterior axis of the cortical subplate, although there were sparse DsRed-labelled fibers within the basomedial (BMA; **Figure 7e–v**) and basolateral (BLA; **Figure 7i–v**) amygdalar nuclei in both Case 1 and Case 7. There was also discrepancy in the relative abundance of DsRed-labeled fibers between the two cases, as Case 1 displayed more fibers in the anterior BMA and BLA (L59–72; **Figure 7h1–o1**; **Table 5**), while Case 7 displayed more fibers in the posterior BMA and BLA (L67–81; **Figure 7l2–u2**; **Table 5**). No fibers were observed in the remaining regions of the CTXsp including the claustrum (CLA; **Figure 7a–m**), endopiriform nucleus (EP; **Figure 7a–v**), lateral amygdala (LA; **Figure 7k–s**), and posterior amygdalar nucleus (PA; **Figure 7r–w**).

#### 3.4.2 Cerebral nuclei

Overall, we observed low-to-moderate amounts of DsRed-labelled fibers across the cerebral nuclei though fibers were more abundant in Case 7 than Case 1 (**Table 5**).

##### Striatum (STR)

The lateral septal complex (LSX) contained a prominent angular fiber band along the border between the caudal (LSc) and rostral (LSr) lateral septum (**Figure 4a**; **Figure 7b–e**). Interestingly, DsRed-labelled fibers in the LSr appeared in punctate in clusters and retained terminal-like morphology. The striatum-like amygdalar nucleus (sAMY) contained few DsRed-labelled within the medial (MEA, **Figure 7i–r**) and central amygdala (CEA, **Figure 7h–q**), however this was not reflected in the intersecting maps (**Table 5**) due to the sparsity of fibers. Similarly, ventral regions within the striatum (STRv), like the nucleus accumbens (ACB; **Figure 7a–c**) contained few DsRed-labelled, however this was also not reflected in the intersecting maps (**Table 5**). We did not observe any fibers in dorsal regions of the striatum (STRd), which comprises the caudoputamen (CP; **Figure 7a–s**).

##### Pallidum (PAL)

Moderate amounts of DsRed-labelled fibers were detected in caudal regions of the pallidum (PALc), particularly ventrally within the posterior bed nucleus of the stria terminalis (BSTp; L56–58; **Figure 4b**; **Figure 7e–g**). Likewise, there were moderate amounts of DsRed-labeled fibers in ventral regions of the pallidum (PALv), which were mainly located in the medial portion of the substantia innominata (SI; e.g., **Figure 7g**) and appeared continuous with that in the amygdalar complex (e.g., **Figure 7j**). Within medial regions of the pallidum (PALm), some fibers were observed within the medial septal nucleus (MS; **Figure 7a–c**) and diagonal band nucleus (NDB; **Figure 7a–f**). Finally, dorsal regions of the pallidum (PALd), including the globulus pallidus (GP; **Figure 7e–o**) contained few fibers, however they did not overlap between Case 1 and Case 7 (**Table 5**).

#### 3.4.3 Thalamus

Within the diencephalon, the thalamus can be subdivided into a collection of regions based on their main functional outputs, including the sensory-motor cortex (DORsm) and polymodal association cortex (DORpm) related thalamus. Overall, DsRed-labelled fibers were more abundant within the DORpm, particularly at midline thalamic regions (e.g., **Figure 7h**).

##### Sensory-motor cortex-related thalamus (DORsm)

The subparafascicular nuclei (SPF; **Figure 7p–x**) comprised DsRed-labelled fibers-of-passage in notable amounts in its magnocellular part (SPFm; **Figure 7p–r**) and moderate amounts ventrally within its parvicellular part (SPFp; **Figure 7s–x**). Likewise, the subparafascicular area (SPA) contained notable ascending projections along the dorsoventral axis (**Figure 7r**). Posteriorly within the DORsm, moderate DsRed-labelled fibers were observed in the peripeduncular nucleus (PP; **Figure 7w–y**). Within individual injection cases, there were only few fibers-of-passage in the ventral group of the dorsal thalamus (VENT; **Table 5**) that includes the ventral medial nucleus (VM; **Figure 7j–r**), ventral anterior-lateral complex (VAL; **Figure 7i–p**), and ventral posterior complex (VP; **Figure 7k–t**) of the thalamus. Sparse DsRed-labelled fibers were in the geniculate group of the dorsal thalamus (GENd), including the medial geniculate complex (MGd; **Figure 7w–y**), and the dorsal part of the lateral geniculate complex (LGd; **Figure 7p–v**).

##### Polymodal association cortex-related thalamus (DORpm)

Regions along the midline of DORpm, such as the midline group (MTN), medial group (MED), and intralaminar nuclei (ILM) of the dorsal thalamus contained abundant DsRed-labelled fibers. The MTN closely neighbours the ZI, and regions like the nucleus of reuniens (RE; **Figure 7f–q**) and paraventricular nucleus of the thalamus (PVT; **Figure 4c**; **Figure 7e–r**) were heavily labeled with DsRed fibers. For instance, the RE contained dense axon terminals that skirted the parataenial nucleus to innervate the anterior PVT (**Figure 4d**; **Figure 7g–i**). Likewise, nuclei in the MED, like the intermediodorsal nucleus of the thalamus (IMD; **Figure 7k–r**), which lies between the PVT and RE, also contained dense DsRed-labelled fibers-of-passage (e.g., **Figure 7n**). Consistently, medial structures within the intralaminar nuclei of the thalamus (ILM) including the central medial (CM; **Figure 7h–q**) and central lateral (CL; **Figure 7k–q**) nucleus of the thalamus contained moderate DsRed-labelled fibers. Although located laterally, the geniculate group of the ventral thalamus (GENv) also contained moderate DsRed-labelled fibers, particularly within the intergeniculate leaflet of the lateral geniculate complex (IGL; **Figure 7s–v**).

Few DsRed-labelled fibers were observed within the lateral (LAT) and anteroventral (ATN) groups of the dorsal thalamus as well as the epithalamus. Noteably, within the LAT the lateral posterior nucleus of the thalamus (LP; **Figure 7n–w**) contained the most DsRed-labelled fibers, particularly within the dorsomedial portion (e.g., **Figure 7r**). The remaining LAT regions such as the posterior complex of the thalamus (PO; **Figure 7l–u**) contained few DsRed-labelled fibers. Majority of the anterior thalamic regions, including the anteroventral (AV; **Figure 7g–k**) and anterodorsal (AD; **Figure 7g–j**) nucleus of the thalamus had sparse DsRed-labelled fibers. Lateral to the AD and AV, DsRed-labeled fibers strikingly surrounded the lateral (**Figure 7i–j** ➔ L62–L63) and dorsal border (**Figure 7k–n**) of the lateral dorsal nucleus of the thalamus (LD). In the epithalamus, comprising the medial (MH; **Figure 7j–s**) and lateral habenula (LH; **Figure 7k–s**), DsRed fibers were sparse, despite the abundance of DsRed fibers in neighboring regions like PVT and CM. Similarly, we observed fre DsRed-labelled fibers in the reticular thalamus (RT; **Figure 7h–r**), despite the RT being in close proximity to the ZI.

#### 3.4.4 Hypothalamus

The diencephalon also includes the hypothalamus, which comprised DsRed-labelled fibers throughout its rostrocaudal extent (**Table 5**).

##### Periventricular zone (PVZ)

The paraventricular hypothalamic nucleus (PVH) contained abundant DsRed-labelled fibers, particularly within the parvocellular (PVHp; **Figure 7e–k**) and magnocellular divisions (PVHm; **Figure 7i–j**). Among regions abutting the third ventricle, DsRed-labelled fibers were observed in the anterior (PVa; **Figure 7h**) and intermediate part (PVi; **Figure 7i–k**) of the periventricular hypothalamic nucleus. However, the arcuate hypothalamic nucleus (ARH; **Figure 7k–q**) contained moderate amounts of DsRed-labelled fibers, particularly at anterior levels (L66–67; **Figure 7k–l**). DsRed-labeled fibers were present in the supraoptic nucleus (SO) of Case 7 (**Figure 7e2–j2**) but largely avoided the SO in Case 1 (**Figure 7e1–j1**).

##### Periventricular region (PVR)

Dense fibers were observed in the neighboring subparaventricular zone (SBPV; **Figure 7i**, **j**) and posteriorly in the dorsomedial hypothalamic nucleus (DMH; **Figure 7l–q**), which was more prominent in Case 7 (**Figure 7l2–q2**). Anteriorly, the median preoptic nucleus (MEPO; **Figure 7b–d**) and medial preoptic area (MPO; **Figure 7b–g**) contained low to moderate DsRed-labelled fibers distributed throughout.

Sparse DsRed-labelled fibers were observed in the ventrolateral (VLPO; **Figure 7d**), anterodorsal (ADP; **Figure 7c–d**), and anteroventral (AVP; **Figure 7c–d**) preoptic nucleus, anteroventral periventricular nucleus (AVPV; **Figure 7c–d**), and suprachiasmatic nucleus (SCH; **Figure 7f–i**). Although individual injection cases displayed moderate DsRed-labelled fibers, the intersecting maps revealed minimal fibers in the posterior part (PVp; **Figure 7r–t**) and preoptic part (PVpo; **Figure 7e–g**) of the PV. Individual injection cases contained sparse DsRed-labelled fibers in the parastrial nucleus (PS; **Figure 7d**), although this was not reflected in the intersecting maps. No DsRed-labelled fibers were observed in the vascular organ of the lamina terminalis (OV; **Figure 7b–c**) and subfornical organ (SFO; **Figure 7g–i**). Unfortunately, we were unable to capture the posterodorsal preoptic nucleus (PD) as we did not collect the level that this region comprises.

##### Medial zone (MEZ)

DsRed-labelled fibers were abundant within regions near the injection site (**Figure 4f**), including the anterior hypothalamic nucleus (AHN; **Figure 4e**; **Figure 7g–k**) and descending division of the PVH (PVHd; **Figure 7i–j**). In posterior regions, abundant DsRed-labelled fibers were observed within the posterior hypothalamus (PH), which appeared to comprise fibers-of-passage travelling towards the periaqueductal gray (PAG; e.g., **Figure 7s–u**), though numerous bouton-like structures were also evident in the PH.

Interestingly, DsRed-labeled fibers largely avoided the ventromedial hypothalamic nucleus (VMH; **Figure 7k–o**), though a few axons may pass through the VMH at posterior levels (L70–73; **Figure 7n–p**). Light to moderate DsRed-labelled fibers were in the ventral (PMv) and dorsal (PMv) premammillary nuclei (**Figure 7r–s**) and even fewer fibers at the ventrolateral border of the mammillary body (MBO; **Figure 7r–w**) and medial preoptic nucleus (MPN).

##### Lateral zone (LZ)

The injection site largely encompassed the LZ, so it was difficult to discern fibers-of-passage from terminal sites of DsRed-labeled fibers. DsRed-labelled fibers in the ZI (**Figure 4f1**; **Figure 7i–v**) were abundant and appeared to travel laterally and arrive at the anterior and caudal parts of the thalamus and midbrain.

DsRed-labelled fibers in the lateral hypothalamic area (LHA) were heterogeneous and varied across *ARA* levels. Majority of DsRed-labelled fibers in the LHA at L67–76 concentrated dorsal to the fornix (**Figure 7l–r**). The anterior LHA (L62–66; **Figure 7i–k**), contained relatively sparse axons, despite anterior regions like the lateral preoptic area (LPO; **Figure 7a–g**), retrochiasmatic area (RCH; **Figure 7i–j**), and tuberal nucleus (TU; **Figure 7k–q**) containing moderate fibers. Posteriorly, there were also moderate fibers within the parasubthalamic (PSTN; **Figure 7p–s**). Few fibers were within the preparasubthalamic (PST; **Figure 7o**) and subthalamic nucleus (STN; **Figure 7o–s**).

##### Median eminence (ME)

DsRed-labelled fibers were detected in the median eminence (ME) of Case 7 (**Figure 7n2–q2**) but not Case 1 (**Figure 7n1–q1**). From our samples, it was unclear whether DsRed-labelled fibers in the ME targeted internal or external lamina.

#### 3.4.5 Midbrain

The midbrain spans L77–113 and can be further subdivided as sensory-related (MBsen), motor related (MBmot), or behavior-state related (MBsta). Unfortunately, we were only able to collect brain slices up to *ARA* L97 for Case 1, thus descriptions of DsRed-labelled fibers posterior to L97 are from Case 7 only. Overall, DsRed-labelled fibers were most abundant within the Mbmot (**Table 5**).

#### Sensory-related midbrain (MBsen)

The MBsen largely comprises the dorsal aspect of the superior colliculus (SC) and inferior colliculus (IC). While DsRed-labeled fibers were relatively low within the dorsal SC including the optic (SCop), superficial gray (SCsg), and zonal layer (SCzo), most fibers were medially distributed within the SCsg (**Figure 4h**; **Figure 7w–ff**; **Figure 8a, b**). The entire IC belonged to the sensory-related part of the midbrain. In the IC, there were moderate but prominent DsRed-labeled fibers in the external nucleus (ICe) and dorsal nucleus (ICd) anteriorly (e.g., **Figure 8b–e**), but DsRed-labelled fibers emerged in the central nucleus (ICc) posteriorly (**Figure 8f**). Abundant DsRed-labelled fibers were also observed in nuclei within the periaqueductal grey area (PAG) such as the medial trigeminal nucleus (MEV; **Figure 8a–c**).

**Figure 8.**
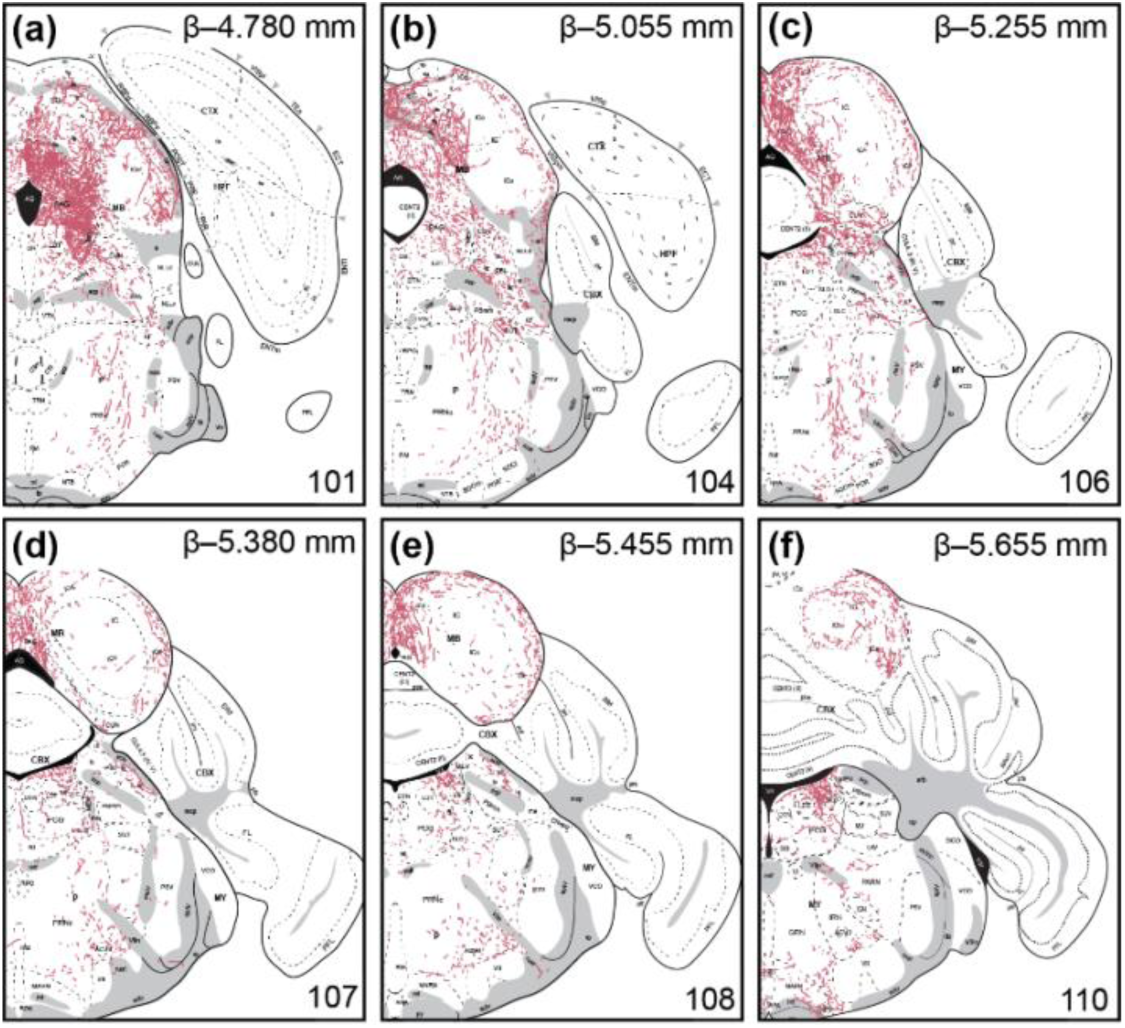
Distribution of fiber projections from *Th-cre* ZI cells in the hindbrain. Mapped distribution of fibers from injection Case 7 with reference to cytoarchitectural boundaries from Nissl-stained tissue using *Allen Reference Atlas* templates (Dong, 2008; **a–f**). Panels include the corresponding atlas level (bottom right), Bregma (*β*; top right), and brain region labels using formal nomenclature from the *ARA*.

There were also low amounts of DsRed-labeled fibers in a collection of brain regions located ventral to the SC and IC that included the nucleus of the brachium (NB; **Figure 7aa**), nucleus sagulum (SAG; **Figure 7ee, ff**) and parabigeminal nucleus (PBG; **Figure 7cc–ff**).

##### Motor-related midbrain (MBmot)

In the dorsal part of the brain, DsRed-labelled fibers were densest and abundant throughout the anteroposterior extent in the PAG and the motor-related SC, but the fiber distribution in these regions was heterogenous. DsRed-labeled fibers concentrated in the dorsal region of the PAG (**Figure 4g**; e.g., **Figure 7z3–ff3**) and along the medial column of the ventral aspects of the SC, especially the deep gray layer (SCdg) and intermediate white layer (SCiw) (**Figure 4h**; **Figure 7v–ff**; **Figure 8a, b**). Ventral to the PAG, midline structures like the oculomotor nucleus (III; e.g., **Figure 7bb**), trochlear nucleus (IV; **Figure 7ff**), red nucleus (RN; e.g., **Figure 7cc**), Edinger-Westphal nucleus (EW; **Figure 7y–z**), and ventral tagmental nucleus (VTN; **Figure 8a–b**) were devoid of DsRed-labeled fibers. Lateral to the PAG, the cuneiform nucleus (CUN) comprised moderate fibers anteriorly (**Figure 8a–d**), while the lateral terminal nucleus of the accessory optic tract (LT; **Figure 8u–v**) was devoid of fibers.

In the ventral part of the brain, DsRed-labled fibers in the midbrain reticular nucleus (MRN; **Figure 7t–ff**, **Figure 8a, b**), including its retrorubral area (RR; **Figure 7z–ee**), were moderate and distributed laterally. Fibers in the ventral tegmental area (VTA; **Figure 7t–z**) were more prominent in the anterior (L81–83; **Figure 7u, v**) than posterior levels (L84–91; **Figure 7w–z**). However, there were no fibers in the reticular part of the substantia nigra (SNr; **Figure 7t–aa**).

In the pretectal region, DsRed-labelled fibers were especially prevalent medially, for example in the medial pretectal area (MPT; **Figure 7s–u**) and less in the nucleus of the posterior commissure (NPC; **Figure 7s–u**), olivary pretectal nucleus (OP; **Figure 7s–u**), or posterior pretectal nucleus (PPT; **Figure 7u**). Other divisions of the pretectal region like the anterior pretectal region (APN; **Figure 7s–w**) and nucleus of the optic tract (NOT; **Figure 7u–w**) were sparsely innervated. Unfortunately, we were unable to capture the anterior tegmental nucleus (AT) as it fell between levels that we did not collect for either injection case.

##### Behavioral state-related midbrain (MBsta)

There were few, if any, DsRed-labelled fibers in the compact part of the substantia nigra (SNc; **Figure 7u–z**), pedunculopontine nucleus (PPN; **Figure 7dd–ff**; **Figure 8a**), or midbrain raphe nuclei (RAmb; **Figure 7v–ff**; **Figure 8a, b**).

#### 3.4.6 Pons

Overall, DsRed-labelled fibers were sparse throughout the pons (**Table 5**).

##### Sensory-related pons (Psen)

There were moderate fibers in the lateral division of the parabrachial nucleus (PBl; **Figure 8b–e**) but less in the medial division (PBm; **Figure 8b–f**). Few DsRed-labelled fibers were present in the nucleus of the lateral leminscus (NLLv; **Figure 7bb–ff**; **Figure 8a, b**), principal sensory nucleus of the trigeminal (PSV; **Figure 8a–f**), and superior olivary complex (SOC; **Figure 7ee–ff**; **Figure 8a–c**). There were no fibers in the Kolliker-Fuse subnucleus (KF; **Figure 8a–d**).

##### Motor-related pons (Pmot)

There were moderate amounts of DsRed-labelled fibers in the pontine central grey (PCG; **Figure 8c–f**) towards the dorsolateral aspect of the region. With the exception of Barrington’s nucleus (B; **Figure 8d–f**), which contained considerable fibers, PCG nuclei such as the dorsal tegmental nucleus (DTN; **Figure 8b–f**) and supragenual nucleus (SG; **Figure 8f**) contained little to no fibers.

There were only few fibers of passage anteriorly in the pontine grey (PG; **Figure 7x–dd**), tegmental reticular nucleus (TRN; **Figure 7z–ff**; **Figure 8a, b**), and supratrigeminal nucleus (SUT; **Figure 8b–e**). There were more fibers laterally in the caudal part of the pontine reticular nucleus (PRNc; **Figure 8b–e**) but largely avoided the motor nucleus of trigeminal (V; **Figure 8a–e**).

##### Behavioral-state related pons (Psta)

This brain division contained very few DsRed-labelled fibers overall. Medial structures like the superior central nucleus raphe (CS; **Figure 7z–ff**; **Figure 8a**), nucleus incertus (NI; **Figure 8c–e**), and nucleus raphe pontis (RPO; **Figure 8b–d**) were particularly devoid of fibers. Laterally, the locus ceruleus (LC; **Figure 8d–f**) and rostral part of the PRN (PRNr; **Figure 7z–ff**; **Figure 8a**) contained a few fibers. Interestingly, while the PAG contained abundant DsRed-labelled fibers, nuclei within the PAG such as the laterodorsal nucleus (LDT; **Figure 8b–f**) did not contain fibers, while the sublaterodorsal nucleus (SLD; **Figure 8b–e**) contained very few. Ventral to the PAG, the subceruleus nucleus (SLC; **Figure 8b–c**) was also devoid of fibers.

#### 3.4.7 Medulla

The medulla spans the remainder of the hindbrain (L97–L132), but our analyses only extended until L110. The majority of DsRed-labelled fibers within the medulla were distributed ventrolaterally and within motor-related regions of the medulla (MYmot; **Table 5**).

##### Sensory-related medulla (MYsen)

No DsRed-labelled fibers were found within MYsen (**Figure 7ff**; **Figure 8a–f**). DsRed-labelled fibers appeared in neighboring ventral regions such as the PRNc in the pons but avoided MY-sen regions like the nucleus of the trapazeoid body (NTB; **Figure 7ff**; **Figure 8a–b**).

##### Motor-related medulla (MYmot)

Majority of fibers are observed within the gigantocellular (GRN), intermediate (IRN), and magnocellular (MARN) reticular nucleus (MARN; **Figure 8f**). Fibers within these regions appeared in the ventrolateral direction and hug the border of the facial motor nucleus (VII), which was devoid of fibers (e.g., **Figure 8f**). A few DsRed-labelled fibers were also seen in the parvicellular reticular thalamus (PARN; **Figure 8f**). Reticular thalamic nuclei such as the gigantocellular (GRN; **Figure 8f**) and intermediate reticular nucleus (IRN; **Figure 8f**) also contained fibers, although sparse.

Sparse DsRed-labelled fibers were observed in the parapyramidal nucleus (PPY; **Figure 8f**) and vestibular nuclei (VNC; **Figure 8f**). No fibers were observed in the abducens nucleus (VI; **Figure 8f**), accessory facial motor nucleus (ACVII; **Figure 8d–f**), and inferior salivary nucleus (ISN; **Figure 78**). Unfortunately, we were unable to collect posterior sections comprising the nucleus ambiguus (AMB), dorsal motor nucleus of the vagus nerve (DMX), infracerebellar nucleus (ICB), inferior olivary complex (IO), linear nucleus of the medulla (LIN), lateral reticula rnucleus (LRN), parasolitary nucleus (PAS), perihypoglossal nuclei (PHY), nucleus x (x), hypoglossal nucleus (XII), or nucleus y (y).

##### Behavioral-state related medulla (MYsta)

None of the midline nuclei, including the nucleus raphe magnus (RM; **Figure 7ff**; **Figure 8a–f**) or nucleus raphe pallidus (RPA; **Figure 8c–f**) in this subdivision contained DsRed-labeled fibers.

#### 3.4.8 Cerebellum

We were unable to capture the entire rostrocaudal extent of the cerebellum from Case 1 or Case 7, but we did not observe DsRed-labelled fibers in the cerebellum in the sections we collected (up to L110; **Figure 8b–f**). However, we were able to confirm the lack of DsRed-labelled fibers in the caudal cerebellum using tissue from other injection cases (e.g., Cases 2–6).

## 4 DISCUSSION

We sought to identify, trace and map the distributions of fibers extending from a population of presumptive DAergic neurons in the mouse ZI that we previously identified to also be GABAergic (Negishi et al., 2020). Interestingly, we found that while almost all ZI *Th-cre* cells expressed *Th* mRNA, only medially distributed ZI *Th-cre* cells co-expressed TH immunoreactivity. We therefore selected injection cases with low (17%) and high (53%) TH coexpression at DsRed-labeled *Th-cre* cells in the medial ZI to assess differences in fiber distribution from cells expressing *Th* mRNA but not TH protein. We mapped and traced brainwide projections from these two injection cases but did not observe stark differences in brain regions labeled, which suggested that TH protein expression minimally affected the distribution of axonal projections from *Th-cre* cells. Given their similarities in projections, we overlapped the mapped fibers from both cases to identify their intersecting points and delineate spatial regions comprising fibers from GABAergic ZI DA cells. We found that DsRed-labelled fibers were distributed throughout the brain but were most prominent in the periaqueductal gray at MBmot and nucleus of reuniens at DORpm.

The ZI has been further subdivided into rostral or caudal and dorsal or ventral components in rats (Kawana and Watanabe, 1981; Romanowski et al., 1985; Nicolelis et al., 1995) and primates (Watson et al., 2014), however there are no clear cytoarchitectural boundaries in the mouse ZI. By validating *Th-cre* expression in the ZI, our findings suggest that the mouse ZI may contain distinct medial and lateral components, as TH-ir cells mostly concentrated within the medial ZI. The mediolateral distribution of TH-ir ZI cells may reflect functional differences, as fibers from medial *Th-cre* ZI cells preferentially targeted midline structures (e.g., lateral septum, nucleus of reuniens, posterior hypothalamic nucleus, periaqueductal gray, superior colliculus), while few fibers were laterally distributed in subcortical (e.g., parvicellular part of subparafascicular nucleus or external nucleus of inferior colliculus) or cortical structures (e.g., central or medial amygdalar nucleus). These projection patterns suggested a medial to lateral conservation of ZI projections, so we may predict that lateral ZI DA cells may preferentially target lateral structures (Kolmac et al., 1998).

The ZI is important for coordinating defensive (Chou et al., 2018; Lin et al., 2023) and hunting behaviors (Shang et al., 2019; Zhao et al., 2019) in rodents. Indeed, DsRed-labelled fibers were abundant in regions that decrease defensive behaviors to promote hunting, and retrograde tracing studies from these regions including the periaqueductal gray (PAG; Kolmac et al., 1998), superior colliculus (SC; Bolton et al., 2015; Montardy et al., 2022), and lateral septum (LS; Wagner et al., 1995) have corroborated our fiber tracing results. Optogenetic stimulation of PAG-innervating ZI GABA cells decreased flight and freezing behaviors (Chou et al., 2018) and increased the motivation to hunt (Zhao et al., 2019). Therefore, ZI GABA cells may promote hunting by suppressing defensive behaviours (Chou et al., 2018) while promoting motivational drive (Zhao et al., 2019). Given the critical role of DA in associative learning during emotionally or physically salient events (Bromberg-Martin et al., 2010), it is possible that projections from ZI Th-cre cells in the PAG could promote learned associations that facilitate hunting. This would be consistent with previously established roles of the dorsolateral PAG that support memory formation and learned association (Di Scala et al., 1987) via aversive stimuli (Di Scala and Sandner, 1989; Castilho and Brandão, 2001). The SC is also an important region for coordinating approach (Shang et al., 2019; Huang et al., 2021) and avoidance behaviours (Cohen and Castro-Alamancos, 2010; Evans et al., 2018). The SC is functionally divided along its mediolateral axis, as the medial SC promotes defence and avoidance and the lateral SC promotes approach and appetitive behaviours (Comoli et al., 2012). Our DsRed-labelled fibers concentrated towards the midline of the SC, therefore ZI Th-cre projections in the SC may inhibit avoidance behaviors. Finally, the LS is a critical region for regulating defensive and aggressive behaviors, as lesions to this area produce “septal rage” (Bard, 1928; Spiegel et al., 1940). Our study revealed dense DsRed-labelled fibers along the border of the dorsolateral LS, which is involved in decreasing fear and freezing responses (Besnard et al., 2019) and preventing the acquisition of fearful memories (Hunsaker et al., 2009; Besnard et al., 2019; Opalka and Wang, 2020). ZI Th-cre cells may thus regulate the acquisition of fearful memories through the LS. Collectively, ZI Th-cre cells densely innervate regions involved in promoting hunting by decreasing defensive behaviors, however the role of DA within these regions is not currently clear.

Dopaminergic ZI cells are a unique subpopulation of ZI GABA cells that project to similar downstream targets, but also innervate additional brain regions. The MBmot comprised the most projection fibers from ZI DA cells, and this was comparable to that by medial ZI GABA cells (Yang et al., 2022). Similarly, both ZI GABA and DA cells do not project to the cerebellum and only sparsely innervate cortical regions like the striatum and pallidum (Yang et al., 2022). By contrast, medial ZI GABA and DA cells can also display differences in their projection targets. Efferent projections from medial ZI GABA cells were abundant within the sensory-motor cortex-related thalamus (DORsm; Yang et al., 2022), but we found that ZI DA cells preferentially projected to the polymodal association cortex-related thalamus (DORpm). Furthermore, ZI GABA cells send strong projections to the pons (Yang et al., 2022), but our ZI DA cells had little or no fibers in the pons. These differences in the distribution of their projections may independent functions of ZI DA cells.

Emerging functions ascribed to ZI DA cells suggested a cooperative relationship with ZI GABA cells to mediate similar outcomes on food- and fear-related behaviors. Optogenetic stimulation of projections from ZI GABA cells in the paraventricular thalamic nucleus (PVT) elicited robust, binge-like feeding and weight gain (Zhang and van den Pol, 2017). Meanwhile, chemogenetic activation of ZI DA cells increases the motivation for feeding by increasing meal frequency without increasing overall food intake (Ye et al., 2023). Since ZI DA cells are also GABAergic (Negishi et al., 2020), it is not known if stimulation elicits both GABA and DA release. However, it stands to reason that independent targeting and stimulation of ZI DA and GABA projections at the PVT would elicit synergistic effects that increase the frequency and amount of food intake. Feeding is also regulated by fear, as a hungry animal would not feed in the presence of a predator. The rostral ZI can modulate anxiety-related behaviors (Zhou et al., 2021), and ZI GABA cells contribute by suppressing anxiety-related escape behaviors (Chou et al., 2018) or fear responses (Venkataraman et al., 2021), while ZI DA cells contribute by encoding learned fear associations (Venkataraman et al., 2021). Collectively, ZI GABA and DA cells innervate the same target sites to coordinate similar behaviors and their coincident outputs may have synergistic outcomes like in feeding and fear association.

### Methodological considerations

Our validation studies revealed a small proportion of ectopic *Egfp* cells that did not express *Th* mRNA or TH-ir. *Th* mRNA levels can decrease in adulthood (Bupesh et al., 2014), thus reduced or trace levels of TH protein may require colchicine pretreatment for immunodetection (Asmus et al., 1992; Asmus and Newman, 1993). Therefore, ectopic *Egfp* cells in the ZI may be related to waning expression of *Th* mRNA in the adult.

*Th* mRNA in the absence of TH-ir has been noted in other brain regions, such as in olfactory bulbs (Min et al., 1994), however the functional relevance of this is still unclear (Savitt et al., 2005). Immunodetection of TH protein may be limited by post-transcriptional silencing, antibody sensitivity, or trace TH concentrations at the soma. We targeted the medial ZI and transduced a higher proportion of TH-ir cells, but the overall connectional profile did not significantly differ from targeting ZI cells only expressing *Th* mRNA. This suggests that ZI cells expressing TH-ir or only *Th* mRNA have similar connectivity profiles. However, future work is required to better understand the developmental trajectory of TH-ir cells in the ZI.

### Conclusion

Overall, medial ZI DA cells have a similar connectivity profile to that of ZI GABA cells, suggesting that ZI DA cells may perform synergistic or opposing functions at the same target sites innervated by ZI GABA cells. Indeed, the studies avaliable demonstrate that ZI DA cells can contribute to nuanced aspects of behaviours previously attributed to ZI GABA cells, including binge-eating, defensive behaviour, associated learning, and hunting. We observed the greatest labelling of ZI DA fibers within motor-related regions of the midbrain, particularly within the dorsal PAG and SC. Future work is required to understand the role of DA within these and other downstream targets.

## Acknowledgments

This work was funded by the National Science Engineering Research Council (NSERC) Discovery Grant RGPIN-2017-06272 (MJC); U. S. National Institutes of Health (NIH) grant SC1GM127251 (AMK); and Howard Hughes Medical Institute education grant for the UTEP PERSIST Brain Mapping & Connectomics Laboratory (AMK). This work was also supported by a NSERC Canadian Graduate Scholarship (CGS) –Master’s (BSB, CDPS, KSB); NSERC CGS – Doctoral (BSB); Internship-Carleton University Research experience for Undergraduate Students (YD); Queen Elizabeth II Graduate Scholarship in Science and Technology (CDSP); Carleton University Work Study Program (TA, MG); U.S. National Science Foundation Grant DUE-1565063 funding the UTEP ACSScellence Program (EM). UTEP PERSIST Brain Mapping & Connectomics Laboratory trainees (MSP, EM) and Eloise E. and Patrick Wieland Fellow (KN) were supported by NIH SC1GM12721 (AMK).

